# Discovery of rare antigen-specific TCRs via replicate profiling

**DOI:** 10.1101/2025.09.30.675572

**Authors:** E.S. Egorov, V.V. Kriukova, I.A. Shagina, G.V. Sharonov, K Lupyr, D.B. Staroverov, D.K. Lukyanov, M.A. Salnikova, R.V. Nikolaev, L. Shelikhova, G. Lopukhova, A. Altunina, M.A. Turchaniniova, O.V. Britanova, E.A. Bryushkova, M.A. Maschan, L.A. Martinez Carrera, N. Jelveh, K. Bisdorf, S.K. Matzke, S. Khorkova, A. Bosio, O. Hardt, A. Franke, D.M. Chudakov, E.O. Serebrovskaya

**Author notes:** Contributed equally (E.S. Egorov, V.V. Kriukova, I.A. Shagina, G.V. Sharonov).

## Abstract

Development of effective vaccines and targeted immunotherapies for cancer, autoimmunity, allergy, and infectious diseases requires comprehensive understanding of functionality and antigenic specificity of involved T cell clones. A major technical challenge remains the high-throughput identification of antigen-specific T cells. Here, we present a rapid cost-efficient TCR discovery assay starting from PBMC that enables ultra-sensitive discovery of clonal alpha-beta paired TCRs responding to individual or pooled peptides. In a small-scale experiment with a single donor, assay identified over 90 SARS-CoV-2-specific CD4^+^ and CD8^+^ TCRβ clonotypes, validated by clonal tracking and comparison against known SARS-CoV-2-specific TCRs. Positioning within the scRNA-Seq map revealed distinct helper T cell subsets involved in primary and secondary response. Further validation in a cohort of five donors identified nearly 1,000 CD4^+^ and CD8^+^ TCR clonotypes specific to viral and fungal peptide antigens. The assay demonstrated exceptional sensitivity in capturing low-frequency clones and allowed accurate TCRα/TCRβ pairing, validated using single-cell transcriptomics. The ability to capture low-frequency antigen-specific TCRs, combined with detailed scRNA-Seq annotation, establishes an integrated pipeline that links antigen-responsive clones to their precise functional phenotypes. This platform provides a robust foundation for dissecting T cell roles in health and disease and accelerates the development of vaccines and immunotherapies.

## Introduction

Functional characterization and transcriptomic mapping of T cells remain incomplete without knowledge of their clonal antigenic specificities. Numerous approaches currently exist to detect T cells specific to target antigens. The most direct method utilizes MHC-multimer staining^1, 2, 3, 4, 5^, which can be coupled with single-cell RNA and TCR sequencing (scRNA-Seq and scTCR-Seq) through oligo-barcoded MHC multimers^6^ or combined with expansion-based pre-enrichment of antigen-specific clones^7^.

Other methods include cultivation-based workflows^8^, assessment of activation markers via flow cytometry and FACS sorting, or detection of cytokine production using intracellular cytokine staining (ICS) flow cytometry^9^. ELISPOT assays can ideally detect approximately 4 cytokine-producing cells per 100,000 peripheral blood mononuclear cells (PBMCs)^10^, but they do not provide information on the T cell receptors (TCRs) of the responding cells. The sensitivity of flow cytometry-based assays is lower, with ICS detecting about 1 in 10,000 PBMCs^11^ and MHC multimer staining detecting about 1 in 5,000 PBMCs^12, 13^.

Here, we relied on the concept of antigen-specific T cell activation or expansion followed by TCR repertoire profiling. This approach—whether applied in bulk^8^ or focused on activated T cells or dividing T cells that progressively dilute cytoplasmic dyes such as carboxyfluorescein succinimidyl ester (CFSE)^14^—enables the identification of T cell clones that are selectively activated and/or proliferate in response to specific antigens and consequently become enriched within the TCR repertoire.

Notably, the random nature of cell sampling, interactions between T cells and potent antigen-presenting cells in culture, varying proliferation potential of individual memory T cells, and high non-specific background substantially limit the applicability of such approaches to relatively large clonal T cell expansions^8^.

To address sampling limitations, we leveraged our prior experience with time-lapse T cell clonal tracking^15, 16^ and identification of locally expanded T cell clones in tumor sections^17^. In both scenarios, we obtained independent biological replicates (at the level of cells) for TCR repertoire profiling. Subsequent bioinformatic analysis identified reproducibly expanded clones with statistical significance, while accounting for sampling noise defined as replicate-to-replicate variability.

Based on this experience and concept, we present a method for identifying low-frequency antigen-specific T cell clones. In the first small-scale experiment, our method detected over 90 clonotypes specific to diverse SARS-COV-2 antigens from a single convalescent donor, with high accuracy confirmed by downstream analyses. Mapping CD4^+^ clones to the scRNA-Seq data revealed their helper T cell subsets, providing a straightforward pipeline for linking T cell clonal specificity to functional programs.

We applied our TCR discovery method to a range of viral and fungal antigens and identified dozens of antigen-specific CD4^+^ and CD8^+^ T cell clones. The methodology demonstrated exceptional sensitivity, capable of detecting clones present at a frequency as low as one per 100,000 T cells.

Furthermore, in 40% of cases we were able to pair TCRα and TCRβ clonotypes based solely on frequency correlation within TCR-Seq data, with accuracy confirmed by scTCR-Seq. This approach enables straightforward *in vitro* validation of identified clones without requiring single-cell transcriptomics.

Overall, we report a robust and highly sensitive approach for identifying antigen-specific memory T cells. This methodology holds considerable promise for accelerating the development of vaccines, immunotherapies, and CAR-T cell therapies by enabling precise characterization of T cell responses and facilitating the engineering of T cell receptors with defined antigen specificities.

## Results

### COVID-specific TCRs discovery

We first focused on searching SARS-CoV2-specific CD4^+^ T cell clones in a donor (D11), who was vaccinated with adenoviral SARS-CoV2 vaccine in Dec2020/Jan2021 and got two registered, PCR-confirmed SARS-CoV2 infections in April 2021 and January 2022 (**Supplementary Note 1, Supplementary Fig. 1, Supplementary Table 1**).

To identify TCR variants responding to SARS-CoV2 antigens for this donor, we implemented and compared three approaches:

### A. TCR discovery via culturing and profiling in replicates

Here, we cultured CFSE-labeled D11 PBMCs in the presence of three distinct mixtures of antigenic peptides, each in three independent replicates. After 7 days, divided (CFSE^low^) CD4^+^ and CD8^+^ T cells were sorted from each replica, yielding 224-7,322 T cells, followed by the sensitive TCR profiling using human TCRαβ RNA Profiling Kit (MiLaboratories/Miltenyi Biotec). TCRβ CDR3 repertoires have been then analyzed with the pipeline based on edgeR, a widely used method to identify differentially expressed genes in RNA-Seq^18^. Here, edgeR algorithm allowed us to identify TCR CDR3 variants that were reproducibly enriched within repertoires of replicates cultured in the presence of the corresponding mixture of antigenic peptides, compared to the other mixtures of antigens (following the same logic as in Ref. 15). Those antigenic peptide pools comprised overlapping 15-mers of 1) SARS-CoV-2 spike protein (S), 2) pooled membrane glycoprotein (M) + nucleoprotein (N), and 3) a mixture of irrelevant control peptides (**Fig. 1**).

EdgeR analysis of obtained repertoires revealed 93 TCRβ clonotypes that specifically and reproducibly proliferated *in vitro* in the presence of S (28 CD4^+^ and 4 CD8^+^ clonotypes) or M+N (59 CD4^+^ and 2 CD8^+^ clonotypes) peptide pools. This bias towards CD4^+^ T cells could be explained by the nature of the 15-mer peptide pools, since the proper processing into shorter 9-10-mer peptides, which are presented by MHC class I to CD8^+^ T cells, may not always occur efficiently—a question that requires further investigation.

**Figure 1.**
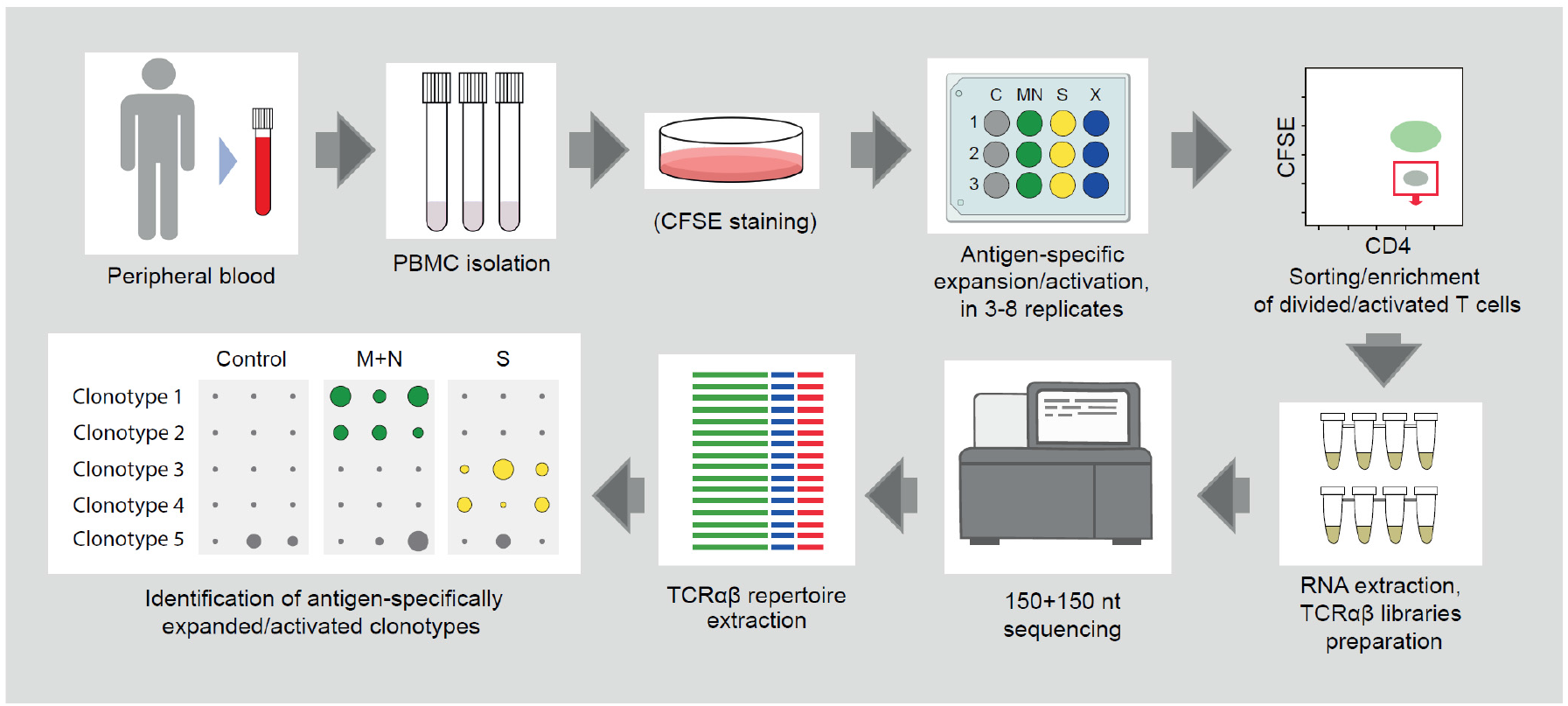
Overview of the TCR discovery concept.

The frequencies of these clonotypes were tracked within the deep peripheral blood TCRβ repertoires of the donor, which were obtained at multiple time points. The analysis revealed that these clonotypes expanded in the donor’s blood exclusively on two time points, which coincided with one or both SARS-CoV-2 infections (**Fig. 2a**), confirming the high accuracy of the method. A group of homologous CD4^+^ Spike-specific TCRβ CDR3s (CASQEGVSNQPQHF, CASSEGASNQPQHF, CASSEGSSSQPQHF, TRBV6-1/TRBJ1-5) was identical or highly homologous to previously described TCRs specific to Spike epitope NLLLQYGSFCTQLNRAL in DRB1^*^15:01 context^19^, also carried by D11. One of the CD4^+^ Spike-specific clonotypes that first appeared after vaccination also responded to the 1st SARS-CoV-2 infection and then dominated in the 2nd SARS-CoV-2 infection, suggesting vaccine-mediated protection (CASSQDSQNLGGSYEQYF, TRBV4-2/TRBJ2-7, **Fig. 2a**).

**Figure 2.**
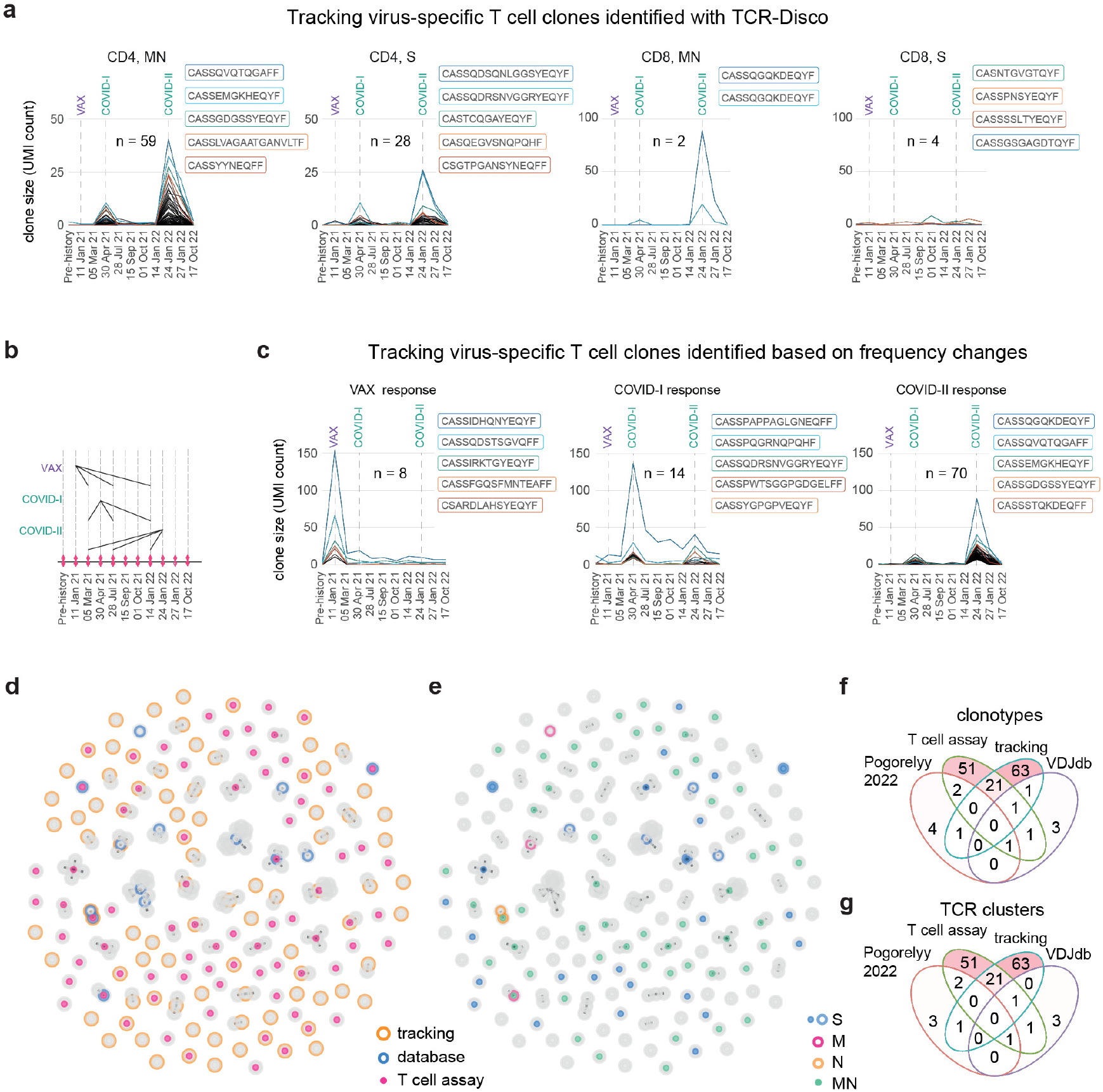
Three methods of capturing COVID-specific TCRβ clonotypes. **a,c**. Longitudinal tracking of TCRβ clonotypes (CDR3nt + V + J) identified with our TCR discovery approach (a) and based on expansion of frequency in target time points (c), using PBMC TCRβ profiling in replicates in sequential time points. Each PBMC TCR-Seq library is downsampled to 45,000 TCRβ encoding cDNA molecules (UMI). TCRβ CDR3 amino acid sequences are shown for the five largest clonotypes on each plot. **b**. Schematic representation of target and reference time points for frequency-based discovery of expanding TCRβ clonotypes. E.g. time point 24 January 2022 was compared to the non-COVID time points 05 March 2021, 28 July 2021, and 14 January 2022. **d**,**e**. TCRβ CDR3 homology clusters identified by the three methods. Each node represents a single CD3aa + V + J clonotype, nodes having one amino acid mismatch in CDR3β are connected with edges. Method (e) and target antigen, if known (f) are shown. **f**,**g**. Venn diagrams representing clonal (g) and CDR3 homology clusters (h) overlap between the methods.

Notably, the frequencies of each of these clonotypes were relatively low, mostly below 0.1% of the whole peripheral repertoire, even at the peak response to the 2nd infection. Their frequency decreased after the resolution of the infection, falling well below the sensitivity of MHC tetramer staining or antigen-specific activation assays^5^. This result demonstrates the potency of our method, which integrates the strengths of antigen-specific T cell enrichment, replicates, and comparative data post-analysis, in identifying low-frequency T cell clones specific to the antigens of interest.

### B. Clonal expansion in corresponding time points

Although our TCR discovery approach allowed us to identify more than 90 SARS-CoV-2-specific TCRβ variants for D11, its results were limited to the three tested antigens, and biased towards CD4^+^ TCRs, thus essentially missing the CD8^+^ TCRs as well as response to non-M/N/S epitopes. To widen our fishing net, we applied a frequency-based approach to identify the TCRβ variants that specifically expanded at the time points of vaccination or SARS-CoV-2 infection in the bulk peripheral TCRβ repertoire of D11, a technique applied previously to reveal vaccine-responsive^15^ or pathogen-specific^16, 20^ T cell clones.

Again, edgeR analysis was applied—now to the bulk peripheral TCRβ repertoires obtained at different time points in biological replicates and normalized to the 45,000 randomly sampled UMI (unique TCRβ cDNA molecules) per replicate per time point. This analysis revealed 8, 14 and 70 TCRβ clonotypes expanding at the time points of vaccination, 1st or 2nd SARS-CoV-2 infections, respectively, compared to the three reference time points (**Fig. 2b,c**). A number of clonotypes that were identified as expanded at the time point of the 1st SARS-CoV-2 infection were also expanding at the 2nd infection, and *vice versa*. At the same time, similar analysis revealed only from 1 to 2 clonal expansions in the control time points compared to the challenge time points (**Supplementary Fig. 2a**). This suggests that most of the edgeR-identified clonotypes that were expanded in peripheral blood in the challenge time points are truly associated with the corresponding challenges.

### C. Screening against known SARS-CoV-2-specific TCRs

To exploit the accumulated knowledge of SARS-CoV-2-specific TCRs with known target peptides and/or restricting human leukocyte antigens (HLAs), we also overlapped peripheral time-tracked TCRβ repertoires of our donor with the VDJdb database^21^ and the dataset of CD4^+^ TCRs enriched in SARS-CoV-2 patients with computationally implied HLA allele specificities^22^. We narrowed our search to the HLA class I and II alleles matching the donor’s HLA (**Supplementary Note 1**). This analysis revealed 7 and 8 clonotypes matching those listed in VDJdb and Pogorelyy *et al*., respectively. The largest expansions of identified clonotypes were observed at time points corresponding to infection with SARS-CoV-2 (**Supplementary Fig. 2b**,**c**).

Finally, we integrated the data on SARS-CoV-2-specific TCRβ clonotypes obtained using the three approaches: 1) antigen-specific culturing; 2) frequency-based capturing; 3) database search (**Supplementary Table 2**). To account for the presence of convergent CDR3 groups^23^ we built single aa-mismatch TCRβ CDR3 homology clusters on the pooled 1)+2)+3) data, visualized on **Fig. 2d,e**. Notably, a number of CDR3 homology clusters and clonotypes were independently supported by 2 or all 3 of the approaches, cross-confirming their specificity (**Fig. 2d-g**).

### Mapping functionality of SARS-CoV-2-responsive T cell clones

To functionally characterize SARS-CoV-2-responsive CD4^+^ cell clones, we performed scRNA-Seq on effector/memory helper T cells at the time points right after COVID-I and COVID-II for the same donor D11. We then mapped D11 scRNA-Seq data onto a larger integrated scRNA-Seq landscape comprising 122 donors^24^, assigning each D11 T cell to its corresponding scRNA-Seq cluster (**Supplementary Fig. 3a**). Next, we highlighted SARS-CoV-2-responsive TCRβ clonotypes identified in the previous section within the scTCR-Seq of D11 (**Fig. 3, Supplementary Fig. 3b**).

**Figure 3.**
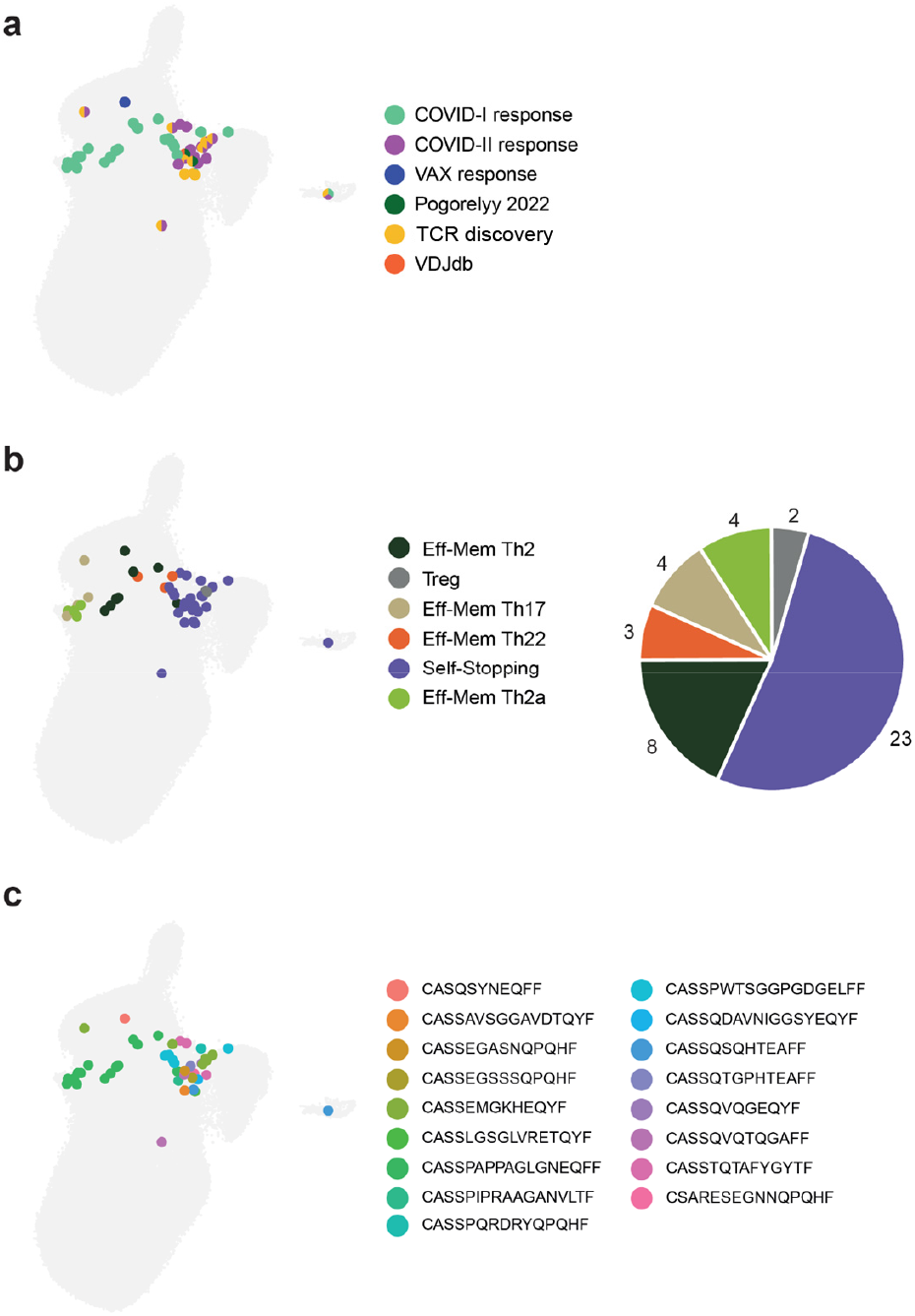
Mapping to the reference dataset shows functional clusters of SARS-CoV-2-specific clones. Positioning of D11 SARS-CoV-2-specific TCRβ CDR3 clonotypes. **a**. SARS-CoV-2-specific cells are colored based on identification method, clonotypes verified by more than one method are shown as small pie charts. **b**. Corresponding scRNA-Seq clusters. Large pie chart shows the proportions of scRNA-Seq clusters in the anti-SARS-CoV-2 response. **c**. TCRβ CDR3 clonotypes.

17 of identified SARS-CoV-2-responsive clonotypes of D11 have been successfully mapped on CD4^+^ T cells localized in several scRNA-Seq clusters. Most of the cells represented “Self-stopping” (*PD-1+, TIGIT+*), Th2, Th2a, and Th17 clusters (**Fig. 3b, Supplementary Fig. 3c**). Most cells recognized as responding to COVID-I based on frequency expansion were also assigned to “Self-stopping”, Th2, Th2a, and Th17 clusters. Most cells responding to the COVID-II as well as identified by antigen-specific culturing assay were assigned to the “Self-stopping” cluster (**Fig. 3a,b, Supplementary Fig. 3c**).

Notably, none of the SARS-CoV-2-responsive clonotypes have been identified among Th1, Temra-Th1, or “Eff-Mem IFN response” clusters. We can suggest that the helper T cell response to COVID-I (moderate illness) was not optimal in the context of viral infection. The presence of “Self-stopping” SARS-CoV-2-specific T cells and their higher clonal heterogeneity could reflect the generally more prominent T cell response to COVID-II (mild illness).

### Further discovery of viral-specific TCR clonotypes

Next, we aimed to extend our study to other viral antigens. In this experiment, similar to the COVID-specific TCR discovery described above, we employed antigen-specific T cell expansion followed by bulk TCR repertoire profiling to identify target antigen-specific TCR clonotypes. However, sorting of CFSE^low^ cell populations was not performed; identification relied solely on the quantitative expansion of target clonotypes in antigen-stimulated conditions. Healthy donor PBMCs were expanded in the presence of different virus-derived CMV, EBV, SARS-CoV-2, and CEF peptide pools. To assess antigen-specific T cell proliferation, pre- and post-cultivation samples were stained with NLV-specific MHC multimer reagent. Median percentage of NLV^+^ CD8^+^ T cells increased to 4.6% in donor B26 and 10.9% in donor C76 after expansion (corresponding to 12.8- and 11-fold expansions, respectively), illustrating strong expansion of the CMV-specific clones (**Fig. 4a,b**). T cell proliferation was also measured by flow cytometry as the percentage of CFSE^low^ CD4^+^ and CD8^+^ T cell subsets. Robust proliferation was detected in samples stimulated with CMV (donors B26 and C76), SARS-CoV-2 (donors C76 and C34), and CEF and EBV (donor C34) peptide pools (**Fig. 4c**).

**Figure 4.**
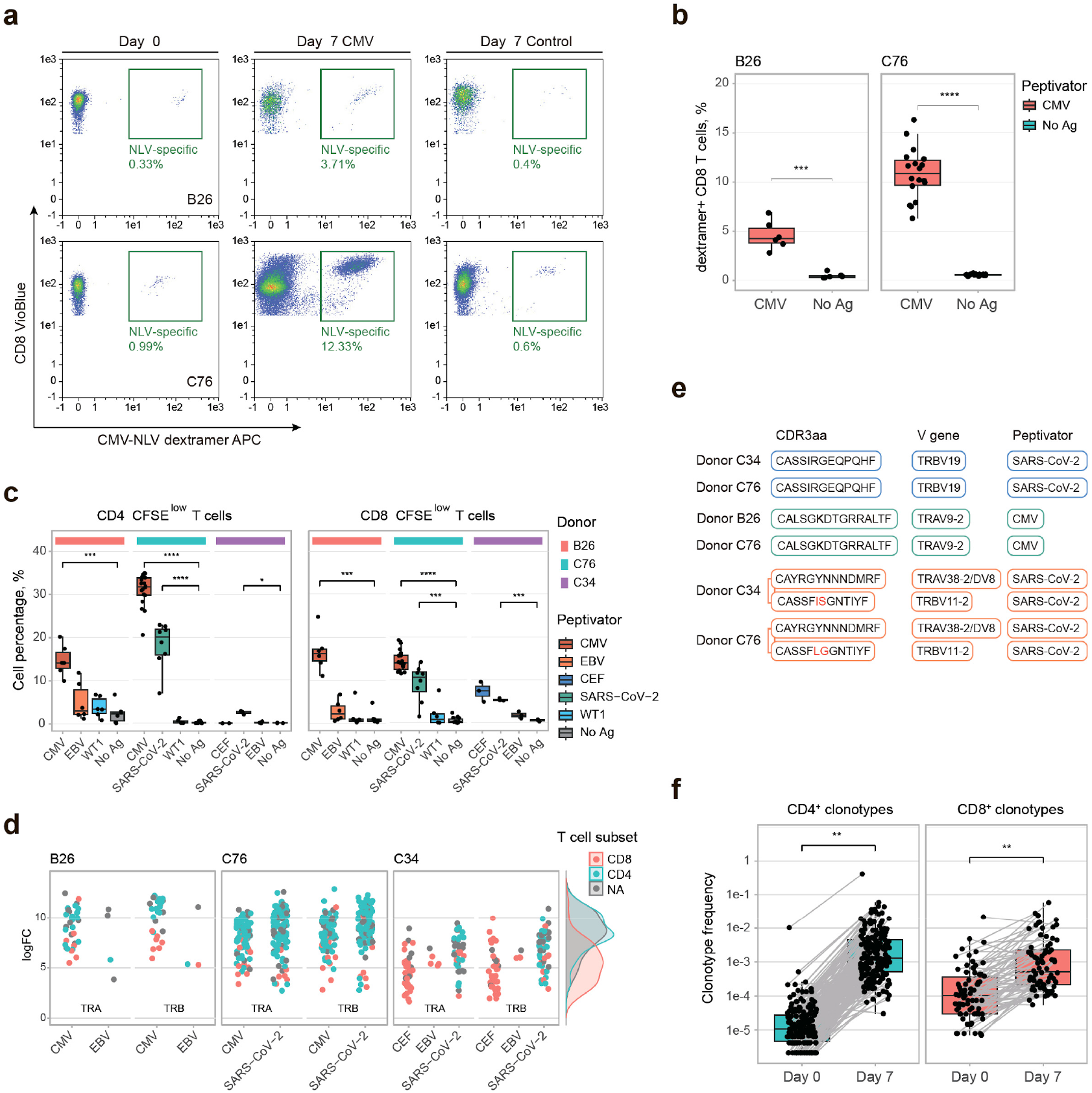
PBMCs culturing with the set of viral peptide pools in three healthy individuals (B26, C34, and C76). **a**. Percentage of CMV-reactive T cells stained by MHC I dextramer (CMV-restricted, NLVPMVATV) before (day 0) and after (day 7) peptide pool stimulation measured by flow cytometry for donors B26 and C76. **b**. Percentage of dextramer positive (NLV^+^) CD8^+^ T cells after 7 days culturing in the presence of the CMV pp65 peptide pool measured by flow cytometry for donors B26 and C76. **c**. Percentage of dividing (CFSE^low^) CD4^+^ and CD8^+^ T cells after 7-day culturing in the presence of the corresponding peptide pool measured by flow cytometry. **d**. TCRα and TCRβ clonotypes specifically expanded upon stimulation with CMV, EBV, CEF, and SARS-CoV-2 peptide pools detected with edgeR pipeline. Each dot represents a unique CDR3 clonotype. edgeR logFC compared to control conditions (all other peptide pools and non-stimulated samples) is shown for each clonotype. **e**. TCRα and TCRβ clonotypes independently detected in several individuals and corresponding predicted paired clonotypes (if available). Predicted clonotype pair CAYRGYNNNDMRF:CASSFISGNTIYF in the donor C34 was confirmed by single-cell TCR-Seq. **f**. Frequencies of antigen-specifically expanded CD4^+^ and CD8^+^ TCRβ clonotypes before and 7 days after stimulation (median value between replicas) in three healthy donors.

TCR discovery with bulk TCR profiling of each cultured replicate identified 579 antigen-specifically expanded TCRα and TCRβ clonotypes (see **Supplementary Fig. 4** for clonotypes identified for donors B26 and C34), with the corresponding edgeR log fold-change (logFC) values plotted in **Fig. 4d**. Several amino acid CDR3 variants were independently detected in two individuals, exhibiting distinct yet convergent nucleotide sequences^25^, demonstrating a public T cell response against common viral epitopes (**Fig. 4e**). To validate our findings, we performed CD137-based magnetic enrichment of antigen-stimulated cells, followed by rapid expansion and single-cell TCR sequencing (IVS workflow), in parallel with expansion-based TCR discovery workflow for one of the donors (C34). SARS-CoV-2-expanded clonotypes detected by TCR discovery workflow were compared with scTCR-Seq data from IVS workflow. Out of 10 most abundant clonotypes found in IVS repertoires, 7 TCRα and 9 TCRβ clonotypes matched with TCR discovery-identified clonotypes. Additionally, we compared the identified antigen-specific clonotypes with the VDJdb database (https://vdjdb.cdr3.net)^21, 26, 27^. Clonotypes with identical CDR3 amino acid sequences, V gene segments, and donor-matched HLA alleles were considered matches. Using this strict rule, we identified multiple matches associated with CMV, EBV, SARS-CoV-2 and InfluenzaA epitopes (summarized in **Supplementary Fig. 5**).

It has been known that long peptides represent a compromise for stimulating both CD4^+^ and CD8^+^ T cell responses^28, 29^. In our experiments, we observed that upon stimulation with CMV pp65, EBV BZLF1, and SARS-CoV-2 Prot_S PepTivators (15-mers with 11 amino acid overlap), antigen-specifically expanded clonotypes were predominantly identified in the CD4^+^ T cell subset. Stimulation with the CEF PepTivator (8–12-mer peptides, known immunodominant epitopes from CMV, EBV, and influenza) identified clones that belonged exclusively in the CD8^+^ T cell subset (**Fig. 4d**).

To assess the sensitivity of our approach, we estimated the initial frequency of identified T cell clones, assuming a yield of approximately 1 UMI per T cell in the non-stimulated PBMC samples. Estimated mean initial frequency was 1 per 10,000 for the CD8^+^ subset and 1 per 100,000 for the CD4^+^ subset (**Fig. 4f**), indicating that our method should allow identification of clones present at frequencies as low as 1-5 T cells per 400,000 PBMCs per replicate. On average, antigen-specific CD4^+^ T cells, which started from lower initial frequencies, expanded more prominently—by about 100-fold—over the 7-day assay, compared to less than 10-fold expansion observed for the initially larger antigen-specific CD8^+^ T cell clones (**Fig. 4f**).

### Computational TCRαβ pairing and pair validation

Although our approach relies on bulk TCR sequencing, which does not preserve the pairing between TCRα and TCRβ clonotypes, we propose a method to predict native TCRαβ pairs for at least some antigen-specifically expanded clonotypes. The underlying idea is that when a T cell proliferates, both the TCRα and TCRβ clonotypes expressed by that cell expand together within the TCR repertoire. Furthermore, if a particular clonotype is completely absent from a sample (which can occur due to the random nature of cell sampling), both the corresponding TCRα and TCRβ clonotypes from that cell would be absent from the repertoire. Therefore, by comparing the frequencies and co-occurrence of expanded TCRα and TCRβ clonotypes across antigen-specific replicates, we can infer native TCRαβ pairs from separate bulk TCRα and TCRβ repertoires. Using this combinatorial correlation method, we predicted 201 TCRαβ clonotype pairs from 977 edgeR-identified TCRα and TCRβ clonotypes (∼40%) (**Supplementary Table 3**).

To validate the accuracy of our pairing method, we compared 14 predicted SARS-CoV-2-specific and 3 predicted EBV-specific TCRαβ pairs from donor C34 with scTCR-Seq data obtained from the IVS experiment for the same donor. Of the 14 predicted SARS-CoV-2 TCRαβ pairs, 10 were confirmed by single-cell (**Fig. 5a,b**).

**Figure 5.**
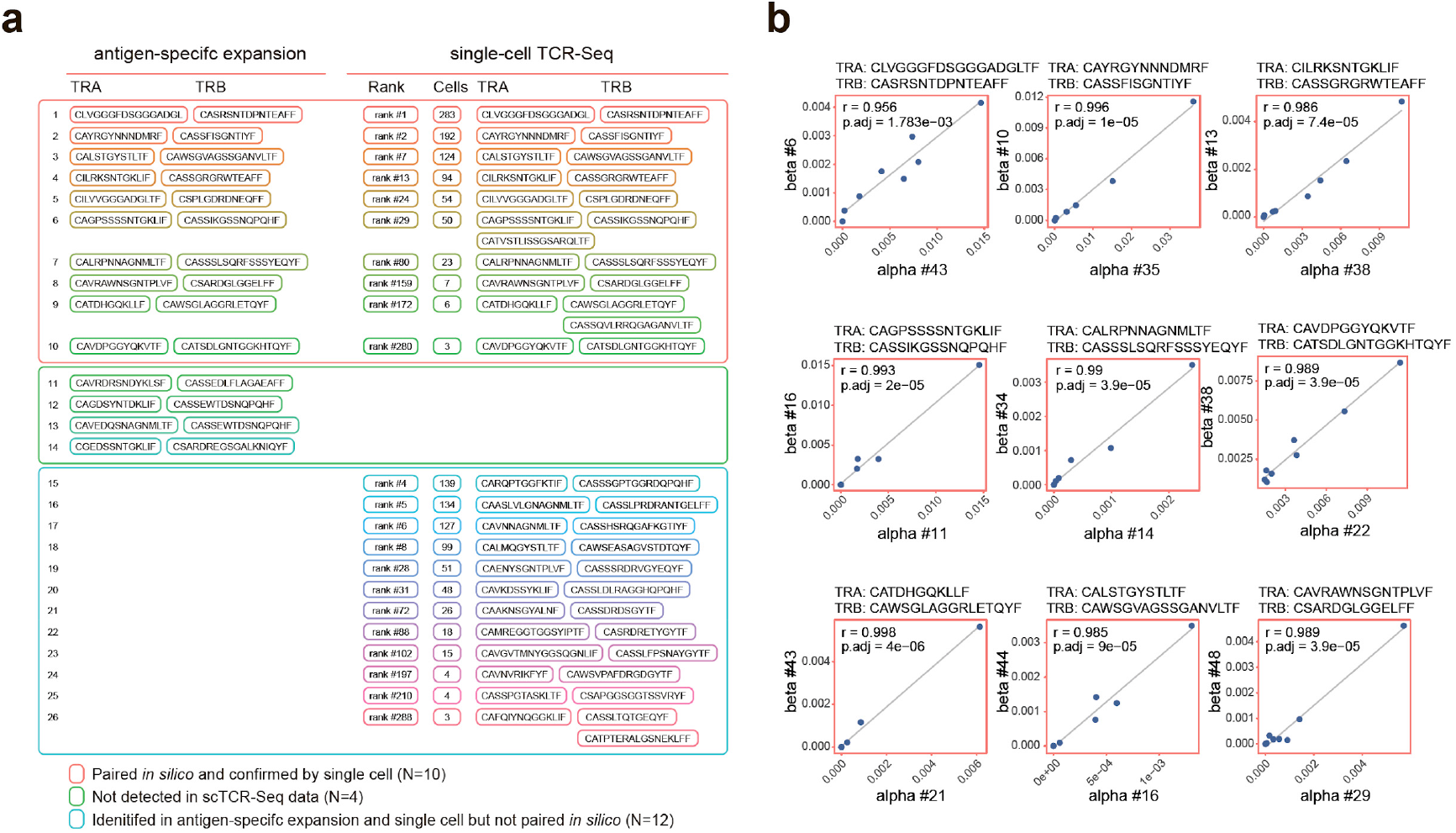
Prediction of TCRαβ clonotype pairs from bulk TCR-Seq data in replicates. **a**. For donor C34, a combinatorial approach predicted 14 SARS-CoV-2-specific clonotype pairs among, 10 pairs confirmed by single cell. Additional 12 SARS-CoV-2-expanded clonotypes were observed in IVS scTCR-Seq data but were not paired *in silico* (bottom part of the table). **b**. Frequency correlation plots for 9 predicted SARS-CoV-2 expanded clonotype pairs that were subsequently confirmed by single cell. Each dot represents the frequency of the predicted TCRαβ clonotype pair (TCRα clonotype - x axis, TCRβ clonotype - y axis) within antigen-specific replicate; r - Pearson correlation coefficient, p.adj – p value after multiple comparison adjustment (Benjamini-Hochberg correction).

The TCRα and TCRβ clonotypes of the remaining 4 pairs were not detected in the scTCR-Seq. Further analysis of the IVS data revealed 12 expanded TCRαβ pairs that were not predicted by our computational method. For EBV, 2 of the 3 predicted pairs were confirmed by single-cell data. Importantly, we observed no mispairing—cases where a TCRα or TCRβ clonotype predicted in one pair was identified in a different pair in the single-cell data—demonstrating the reliability of our pairing approach.

VDJdb includes paired TCRαβ clonotypes and can also be used for pairing validation. Searching the database, we found a match for the SARS-CoV-2-specific clone from donor C76 with the predicted TCRαβ pair CAVHTSGTYKYIF_TRAV21 / CASSLEDTNYGYTF_TRBV7-2, corresponding to a known clone specific for the Spike epitope NQKLIANQF presented by HLA-B^*^15, an allele confirmed in this donor.

Moreover, the TCRα clonotype CAYRGYNNNDMRF_TRAV38-2DV8 from donor C34, whose predicted pairing with TCRβ CASSF**IS**GNTIYF_TRBV11-2 was confirmed by single-cell sequencing, was also independently detected in donor C76 in the context of SARS-CoV-2 response (**Fig. 4e**). Notably, the corresponding TCRα clonotype in donor C76 has a highly homologous predicted TCRβ partner, CASSF**LG**GNTIYF_TRBV11-2. This finding represents a convergent response against SARS-CoV-2 Spike epitope and further demonstrates high reproducibility and effectiveness of our *in silico* pairing method.

### Discovery of fungus-specific TCRs

After successfully identifying several dozen viral-reactive TCRs, we decided to challenge our workflow with potentially weaker stimuli. As an appropriate model we focused on TCR response against two fungal pathogens, *Candida albicans* and *Aspergillus fumigatus*. Despite their distinct biology and infection mechanisms, both pathogens can cause serious health complications, particularly in immunocompromised individuals, including patients undergoing chemotherapy, organ transplant recipients, and those receiving immunosuppressive medications. Understanding the features of TCR response to these pathogens could be crucial for managing infections and improving clinical outcomes in affected patients.

PBMCs from the healthy donors pre-screened for anti-fungal T cell response were stained with CFSE and stimulated with MP65 (*C*.*albicans*) and F22 (*A*.*fumigatus*) peptide pools. PBMCs of one donor were additionally stimulated with Catalase B peptide pool (*A*.*fumigatus*). For all 3 donors, EBV consensus peptide pool was used as a positive control, and wells without antigen served as non-stimulated controls. PBMCs were cultured for 7 days in eight individual replicates per condition. Flow cytometry analysis of proliferating (CFSE^low^) CD4^+^ and CD8^+^ subsets after cultivation confirmed T cell response against fungal peptides in two donors (**Fig. 6a**). The third donor responded only to EBV peptides in the CD8^+^ T cell subset (data not shown) and was excluded from subsequent TCR library generation. Cultured replicates from the two responding donors were lyzed in RLT buffer and processed for bulk TCR library preparation and TCR repertoire profiling as described above.

**Figure 6.**
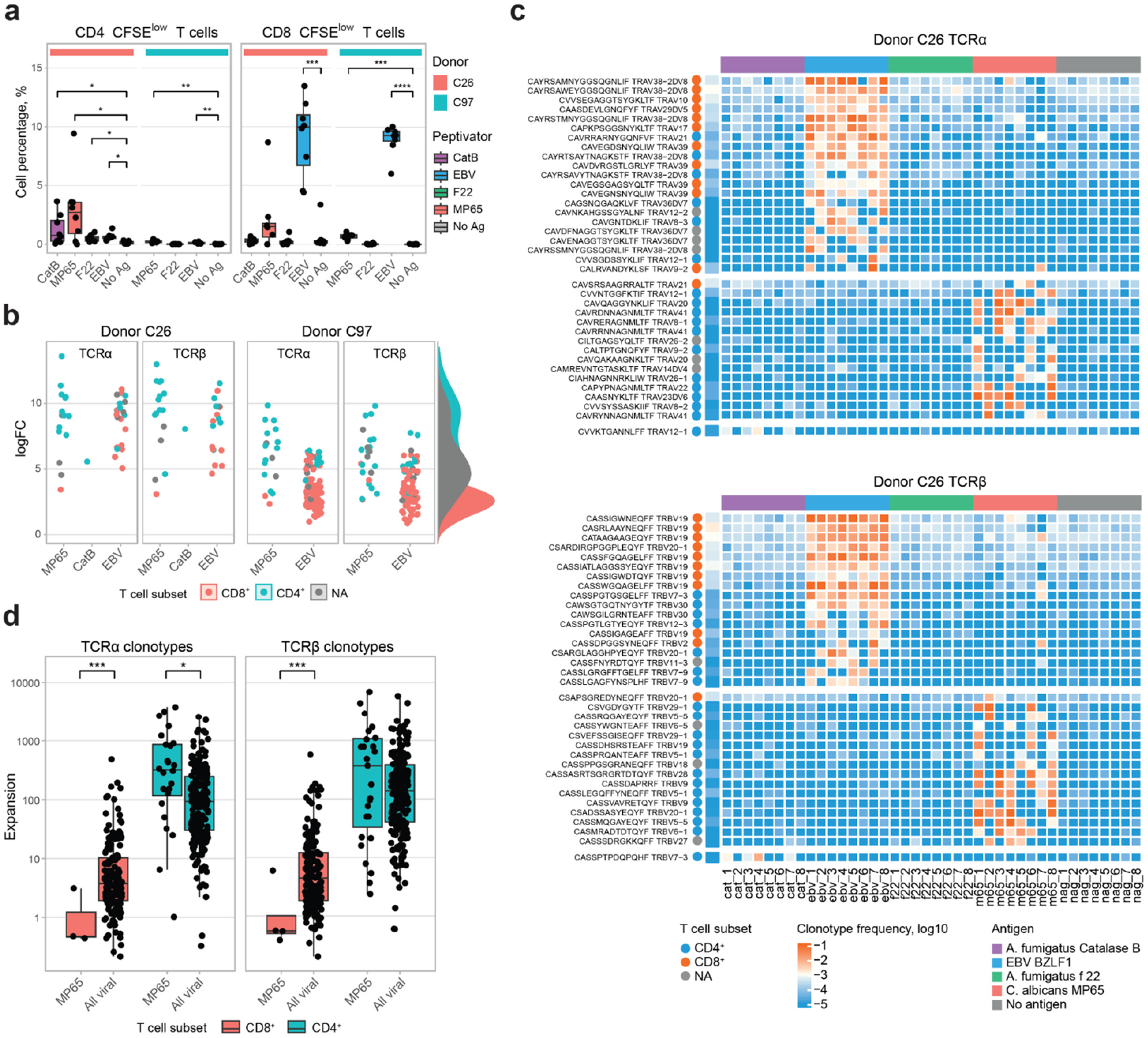
PBMCs stimulation with fungal peptide pools, healthy donors. **a**. Percentage of dividing (CFSE^low^) CD4^+^ and CD8^+^ T cells after 7 days culturing in the presence of the corresponding peptide pool measured by flow cytometry. **b**. TCRα and TCRβ clonotypes expanded upon stimulation with MP65, Catalase B, and EBV peptide pools, identified using the edgeR pipeline. **c**. Frequency of TCRα and TCRβ clonotypes specifically expanded in response to C. albicans MP65, A. fumigatus Catalase B, and EBV BZLF1 PepTivators, donor C26. **d**. Expansion levels of TCRα and TCRβ clonotypes following stimulation with the MP65 peptide pool (donors C26 and C97) and with different viral peptide pools (CMV, CEF, EBV, and SARS-CoV-2; summarized data from the donors B26, C34, C76 and EBV-specific clonotypes from donors C26 and C97).

Using the established workflow, we identified 18 TCRα clonotypes in donor C97, of which 13 were assigned as CD4^+^ and 2 as CD8^+^, and 23 TCRβ clonotypes, with 15 assigned CD4^+^ and 3 as CD8^+^, as reliably expanded upon MP65 peptide pool stimulation (**Fig. 6b, Supplementary 6**). For the donor C26, we identified reliably expanded clonotypes upon MP65 and F22 peptide pools stimulations: for MP65, 15 TCRα clonotypes (11 CD4^+^, 1 CD8^+^) and 16 TCRβ clonotypes (12 CD4^+^, 1 CD8^+^); for Catalase B, 1 TCRα CD4^+^ and 1 TCRβ CD4^+^ clonotype (**Fig. 6b,c**). As with the CMV and SARS-CoV-2 experiments, the response to fungal antigens was predominantly observed in the CD4^+^ T cell subset, likely due to the peptide pools primarily consisting of 15-mer sequences (**Fig. 6b-d**).

## Discussion

Technologies for activation/expansion and enrichment of antigen-specific T lymphocytes have been successfully developed over the past decades. At the same time, despite its great potential, the power of bioinformatic analysis of T cell receptor repertoires to identify antigen-specific clonotypes amidst the noise of non-specific activation and proliferation remains underutilized. Here we report cost-efficient and sensitive methodolgy for identifying antigen-specific T cell clones in patient blood samples. Our method employs classical approaches of antigen-specific cultivation and optionally enrichment of activated/proliferated T lymphocytes^8, 14^, but is further empowered by biological replicates and statistical analysis of obtained TCR repertoires. We demonstrate that antigen-specific clones can be identified with high sensitivity—down to a single T cell per replicate—as well as with high reliability.

Our TCR discovery approach identified more than 80 CD4^+^ SARS-CoV2-specific T cell clones in a single donor using a simple, short assay. Expansion of these clones in longitudinal TCR repertoire-based tracking ideally coincided with two documented SARS-CoV2 infections of the donor, confirming accuracy of the method. We further mapped SARS-CoV2-specific T cell clones to the scRNA-Seq data of the donor, integrated and annotated within the global CD4^+^ T cell phenotype reference map. This enabled us to determine the functional scRNA-Seq clusters to which these clones belonged and, consequently, to infer the nature of the patient’s immune response to the first and second infections. We next expanded this approach to encompass diverse viral antigens, identifying nearly 600 antigen-specific TCRα and TCRβ CDR3s from three donors. These findings were further validated using CD137-based enrichment and mapping to the VDJdb database. Additionally, to assess method ability to identify TCRs specific to non-viral, potentially weaker antigenic stimuli, we searched for fungi-specific T cells in healthy donors and successfully identified dozens of antigen-specific TCRs.

### Sensitivity considerations and limitations

The initial PBMC frequency of identified antigen-specific clones was estimated as 1 per 10,000 T cells for the CD8^+^ subset and 1 per 100,000 T cells for the CD4^+^ subset (**Fig. 4f**). This demonstrates the high sensitivity of our method, enabling the detection of antigen-specific clones present at frequencies as low as 1–5 T cells per 400,000 PBMC replicate. A remaining limitation of our approach is that it targets only antigen-experienced memory T cell clones, which are pre-expanded and represented in the sample by at least several cells. Only under these conditions can we utilize the replicate approach, which requires that the clone is detected in multiple individual wells. This implies that naive T cells—which are very diverse in terms of their TCR identity, and are essentially singletons—will not be detected by our discovery algorithm, as they are unlikely to appear in more than one individual culture. To approach naive T cells, we would need to pre-expand the sample in the presence of the antigen of interest, let it rest until the cell division subsides, and then perform the expansion-based discovery experiment.*Specificity considerations*. Additionally to edgeR statistics, to ensure specificity of identification of antigen-specific TCR chains, we set a certain threshold for the identification in replicates. For experiments with eight replicates, we required detection in at least three out of eight replicates, based on the following rationale. The average frequency of expanded clonotypes within the antigen-specific sample was approximately 1 × 10^−3^, for both CD4^+^ and CD8^+^ T cells. Consequently, the probability of detecting an expanded/activated clonotype in 3 out of 8 target replicates by chance is (1 × 10^−3^)^3^ = 1 × 10^−9^. This indicates that such an observation is unlikely to result from random, non-specific T cell proliferation. Nevertheless, this threshold is somewhat arbitrary, and may be modified depending on the aim of the experiment (sensitivity *versus* specificity).

### CD4^+^ versus CD8^+^ TCR discovery

The successful discovery of antigen-specific TCRs was strongly associated with peptide length: stimulation with long 15-mer peptides preferentially lead to CD4^+^ TCR clonotype discovery, whereas with shorter 9-12aa peptide stimulation, we preferentially identified CD8^+^ TCR clonotypes, consistent with previous studies^28, 29^. According to repertoire analysis, antigen-specific CD4^+^ T cells expanded much more prominently—by approximately 100-fold over a 7-day assay— compared to less than 10-fold expansion of antigen-specific CD8^+^ T cell clones. Considering that the mean initial frequency of identified CD8^+^ clonotypes was 10 times higher than that of CD4^+^ cells (**Fig. 4f**), we speculate that due to higher initial clonality of CD8^+^ repertoire, CD8^+^ T cells have less “room for expansion” before they reach a plateau defined by dynamic competition for nutritional substances and/or cytokines etc. This view is supported by the observation that both CD4^+^ and CD8^+^ T cells identified as antigen-specifically expanding reach approximately the same post-expansion frequency after 7 days of cultivation.

### TCRα/TCRβ pairing

Stochastic nature of antigen-specific T cell sampling and proliferation naturally leads to variability in the frequency of each expanded clone in different replicates. This allowed us to employ frequency correlation-based pairing of TCRα and TCRβ chains^30, 31, 32^. With this approach we could identify 201 TCRαβ clonotype pairs from 977 edgeR-identified TCRα and TCRβ clonotypes, and high accuracy of pairing was confirmed by scTCR-seq data, VDJdb analysis, and presence of highly similar SARS-CoV-2 Spike-specific paired TCRαβ clonotypes in two donors.

We report highly sensitive, cost-efficient, and easy-to-use methodology to identify antigen-specific TCRs in a 10-day assay that starts directly from PBMCs and requires no labor-intensive procedures. This approach should be widely applicable in studies of cancer, autoimmunity, infections, and vaccinations, serving as a convenient, reliable, and sensitive tool for the rapid identification of antigen-specific effector and memory T cell clones.

## Data availability

Raw sequencing data is available in NCBI Sequence Read Archive (BioProject: PRJNA995237). Processed Seurat objects with CD4^+^ T cell reference and scRNA-Seq datasets from D11 are available for download at https://figshare.com/projects/T_helper_subsets_Kriukova_et_al_/173466. PBMC ds45k dataset (D11, TCRβ repertoires from PBMCs in replicates, downsampled to 45,000 UMI in each sample) is deposited in our GitHub repository (Th_kriukova/outs/ds45k) at https://github.com/kriukovav/Th_kriukova.

## Code availability

Code for analysis (TCRβ repertoires from PBMCs) and all figures (scRNA-Seq and TCRβ repertoires) is available at https://github.com/kriukovav/Th_kriukova. R package wrapping Seurat^33^ functions to perform reference mapping to CD4^+^ T cell scRNA-Seq dataset is available at https://github.com/kriukovav/CD4map.

## Acknowledgements

We thank Dr. Imran Siddiqi, Dr. Andrzej Dzionek and Dr. Ian Hardy (all from Miltenyi Biotec) for their help in the implementation of IVS protocol. We thank Vadim Karnaukhov, Denis Syrko, Viktor Kotlyar, and Dmitry Bolotin for assistance with data analysis, and Maria Vakhitova, Tatiana Grigorieva, Ilgar Mamedov, and the NGS Laboratory (IKMB, Kiel University) for assistance with sequencing.

## Funding

Supported by RSF grant №25-75-30013. Sequencing was supported by the DFG Research Infrastructure NGS_CC (project 407495230) as part of the Next Generation Sequencing Competence Network (project 423957469).

## Ethics statement

For donors B26, C76, C26, C97 and C34, 20-100 ml of whole blood were taken from healthy volunteers within the internal blood donation program of Miltenyi Biotec after informed consent. The blood donation program was approved by a positive vote with the procedure number 2020272 from the ethical committee of the “Ärztekammer Nordrhein” from October 29, 2020. For donor D11, ethical approval was granted by the Institutional Review Board of the Pirogov Russian National Research Medical University. The study was conducted in accordance with the Declaration of Helsinki.

## Declaration of competing interests

The authors declare the following financial interests/personal relationships which may be considered as potential competing interests: the authors Evgeniy S. Egorov, Lilian Andrea Martinez Carrera, Nojan Jelveh, Kevin Bisdorf, Simona Karolin Matzke, Svetlana Khorkova, Andreas Bosio, Olaf Hardt, and Ekaterina O. Serebrovskaya are employees of Miltenyi Biotec B.V. & Co. KG. Other authors declare they have no competing interests.

## Author contributions statement

EES, KVV, SIA, SGV, LK, SDB, LDK, SMA, NRV, SL, LG, AA, TMA, BOV, BEA, MLA, JN, BK, MSK, KS - experimental work and data analysis. EES, KVV, LK, KS, MM, BA, HO, FA, SEO, CDM - manuscript preparation. KS, MM, BA, HO, FA, SEO, CDM - supervision. CDM - concept.

## Online Methods

### IFN-γ ELISpot

PBMCs were isolated via standard density gradient centrifugation, quantified using a LUNA-II Automated Cell Counter (Logos Biosystems), and resuspended at 3 × 10^6^ cells/mL in warm (37°C) CTL-Test™ Medium (Cellular Technology Ltd.) containing 1% L-glutamine (Thermo Fisher Scientific). Cells were then plated in 96-well anti-IFN-γ-coated ELISpot plates (Human IFN-γ Single-Color ELISPOT Kit, Cellular Technology Ltd.) at 3 × 10^5^ cells/well in the presence of recombinant SARS-CoV-2 nucleocapsid phosphoprotein (N) or recombinant SARS-CoV-2 spike glycoprotein (S) in duplicate (each 1 µg/mL, Miltenyi Biotec). Negative control wells lacked viral proteins, and positive control wells contained phytohemagglutinin (2.5 ng/mL, Sigma-Aldrich). Plates were incubated for 18 h at 37°C in the presence of 5% CO^2^. Assays were developed according to the manufacturer’s instructions, and spots were counted using an ImmunoSpot S6 Universal Analyzer (Cellular Technology Ltd.).

### TCR discovery assay

PBMCs were freshly isolated with density gradient, stained with CFSE (450 nM) and seeded in complete RPMI complete (RPMI 1640 (PANECO) + 10% heat-inactivated fetal bovine serum + 20 ng/mL IL21(SciStore)) or RP5 (RPMI 1640 (Biowest) + 5% heat-inactivated human serum (Capricorn) + 2 mM L-glutamine (Lonza) + NEAA (MP Biomedicals) + Pen/Strep + pyruvate (Biowest) + 50 ng/ml IL21 (Miltenyi Biotec)). The experiment included 3 - 5 target antigen conditions, and control stimulation either with a pool of irrelevant peptides (all at a final concentration of 0.6 nmol/peptide/mL) or without any antigen (see **Table 1** for conditions used for specific experiments). Cells were stained with anti-CD3, anti-CD8, and/or anti-CD4 antibodies (see **Table 1**) on day 6-7. In some experiments, CFSE^low^CD4^+^ and CFSE^low^CD8^+^ T cells were sorted directly into RLT buffer (200 µL, QIAGEN) using a FACSAria III (BD Biosciences). In other experiments, the plate was centrifuged for 5min at 350g, supernatant was discarded by pipetting, then the cell pellet was lyzed in 200ul RLT buffer (Qiagen) and stored at -70 °C. Additionally, for no-CFSE-sorting experiments, two aliquots of 2 ^*^ 10^6^ PBMCs each were taken for either CD4^+^ or CD8^+^ T cell enrichment with CD4^+^ and CD8^+^ MicroBeads (Miltenyi Biotec), according to the manufacturer’s protocol. Positive fractions of CD4^+^ and CD8^+^ T cells after elution were also lyzed in 200 uL RLT buffer (Qiagen) and stored at -70 °C. Then, RNA was isolated using RNeasy Micro Kit (>500 cells/sample, QIAGEN) or with TRIzol Reagent (<500 cells/sample, Thermo Fisher Scientific).

**Table 1.** Experimental layout: antigens, replicates, antibodies, donors.

| parameter | COVID-specific TCRs discovery | Discovery of viral-specific TCR clonotypes |  |  | Discovery of fungus-specific TCR clonotypes |  | IVS |
| --- | --- | --- | --- | --- | --- | --- | --- |
| donor ID | D1 | B26 | C34 | C76 | C26 | C97 | C34 |
| Seeding conditions | 0.5 x 10<sup>6</sup> cells / well, 48-well plate | 0.2 x 10<sup>6</sup> cells / well, 96-well plate |  |  |  |  |  |

| CFSE-based sorting |  |  | + | - | - | - | - | - | - |
| --- | --- | --- | --- | --- | --- | --- | --- | --- | --- |
| medium |  |  | RPMI complete | RP5 |  |  |  |  |  |
| conditions |  |  | 3 | 4 | 4 | 4 | 5 | 4 | 2 |
| replicates |  |  | 3 | 8 | 8 | 8 | 8 | 8 | 2 |
| antigens<br>(PepTivator<br>s, Miltenyi<br>Biotec) | SARS-CoV-2<br>Prot_S | 130-126-701 | + | - | + | + | - | - | + |
|  | SARS-CoV-2<br>Prot_M and<br>SARS-CoV-2<br>Prot_N | 130-126-698,<br>130-126-702 | + | - | - | - | - | - | - |
|  | irrelevant peptide<br>mix |  | + | - | - | - | - | - | - |
|  | CMV pp65 | 130-093-438 | - | + | - | + | - | - | - |
|  | EBV consensus | 130-103-462 | - | - | - | - | + | + | - |
|  | EBV BZLF1 | 130-093-612 | - | + | + | - | - | - | + |
|  | WT1 | 130-095-916 | - | + | - | + | - | - | - |
|  | CEF MHC Class<br>I Plus | 130-098-426 | - | - | + | - | - | - | - |
|  | C.albicans MP65 | 130-096-776 | - | - | - | - | + | + | - |
|  | A. fumigatus F22 | 130-099-776 | - | - | - | - | + | + | - |
|  | A. fumigatus<br>catalase B | 130-097-291 | - | - | - | - | + | - | - |
| antibodies | anti-CD4–Alexa<br>Fluor 647 (clone<br>RPA-T4,<br>BioLegend) | 300520 | + |  |  |  |  |  |  |
|  | CD8–BV421<br>(clone SK1,<br>BioLegend) | 344748 | + |  |  |  |  |  |  |
|  | anti-CD3-PEVio770, clone<br>BW264/56,<br>Miltenyi Biotec | 130-113-130 |  | + | + | + | + | + | + |
|  | anti-CD8-VioBlue, clone<br>REA734, Miltenyi<br>Biotec | 130-110-683 |  | + | + | + | + | + | + |
|  | antiCD137, clone<br>REA765, Miltenyi<br>Biotec | 130-110-764 |  |  |  |  |  |  | + |
|  | CliniMACS<br>CD137 Biotin | 220-001-830 |  |  |  |  |  |  | + |
| Microbeads<br>(Miltenyi<br>Biotec) | CliniMACS anti-Biotin MB | 200-070-120 |  |  |  |  |  |  | + |
|  | CD4+MicroBead<br>s | 130-045-101 |  | + | + | + | + | + |  |
|  | CD8+MicroBead<br>s | 130-045-201 |  | + | + | + | + | + |  |
| Other<br>staining<br>reagents | HLA-A*0201<br>NLVPMVATV<br>pp65 CMV APC<br>Dextramer,<br>Immudex | WB02132 |  | + |  | + |  |  |  |

TCR libraries were constructed using the human TCRαβ RNA Profiling Kit (MiLaboratories/Miltenyi Biotec). Sequencing was performed on Illumina platform, 150+150 bp paired reads, at least 50 reads per input cell, and raw data analysis were performed as described below. Significantly expanded clonotypes were identified using EdgeR library^18, 34, 35^. UMI counts were filtered using the filterByExpr function with the following parameters: min.count = 1, min.total.count = 5, large.n = 1, and min.prop = 0.5. Count normalization and estimated dispersion by the default method were performed using the trimmed mean of M values (TMM). Each group was tested against all others using a quasi-likelihood (QL) negative binomial generalized log-linear model followed by the F-test. Final clonotypes had an FDR <0.05 and a logFC ≥4. The code is available as part of the TCRgrapher library at https://github.com/KseniaMIPT/tcrgrapher (edgeR_pipeline function) ^36^.

### *In vitro* stimulation and CD137-based enrichment of PBMCs (IVS)

Freshly isolated PBMCs were resuspended in RPMI-1640 medium supplemented with 5% human AB serum (Lonza, Basel, Switzerland) and 2 mM L-glutamine (GE Healthcare). Subsequently, 1 × 108 PBMCs were stimulated at 1 × 10^7^ cells/ml with the BZLF1 and SARS-Cov-2_Prot_S PepTivators (Miltenyi Biotec); each peptide was present at a final concentration of 0·6 nmol/ml. After 45 hours of stimulation, cells were labelled with CliniMACS CD137 GMP Biotin antibody for 15 min, washed and then incubated with CliniMACS Anti-Biotin Reagent for 30 min (Miltenyi Biotec). All incubations were carried out at RT with continuous rotation using the MACSmix instrument (Miltenyi Biotec). After the final washing step, cells were magnetically captured on LS columns, attached to QuadroMACS separator (both from Miltenyi Biotec). Positive and negative fractions were then seeded together with gamma-irradiated feeder cells in TexMACS medium (Miltenyi Biotec) with 5% human AB serum (Capricorn), penicillin/streptomycin, 30 ng/ml MACS GMP OKT3 antibody and 3000 IU/ml MACS GMP IL2 (both from Miltenyi Biotec). Cells were expanded for 14 days, with media exchange at days 5, 7, 9 and 11 without addition of the OKT3 antibody. On day 14, cells were harvested and processed for scTCR-seq.

### Flow cytometry

Dextramer staining was carried out according to the protocol provided by the manufacturer (Immudex). Shortly, uncultured or cultured PBMCs from donors B26 and C76 were resuspended in 50ul PEB buffer (PBS with 1mM EDTA and 0.5% BSA, pH7.2) and incubated with 10ul NLV Dextramer reagent (WB02132, Immudex) for 10 min at RT in the dark. Then, cells were counterstained with CD3-PEVio770 and CD8-VioBlue (**Table 1**) for another 20 min on RT in the dark. After incubation, cells were washed twice with 1.5 ml PEB, resuspended in 500 ul PEB buffer and then acquired on MACSQuant 16 instrument (Miltenyi Biotec). Lymphocytes were initially gated using forward versus side scatter, then by CD3 and CD8 labelling. CD3^+^CD8^-^ cells were considered CD4^+^. CFSE staining and dextramer staining was analyzed separately for CD4^+^ and CD8^+^ cell populations.

### RNA isolation

Bulk RNA were isolated fromCD4^+^/CD8^+^ pre-cultivation samples and samples after 7 days culturing with corresponding PepTivator, lyzed in RLT buffer, using RNeasy Micro kit (Qiagen) and RNeasy Mini Spin columns (Qiagen) according to manufacturer protocol excluding DNAse treatment step to prevent additional material loss. For TCR repertoire profiling ½ volume of initial RLT sample was used. The second part of RLT samples was frozen back and stored at -70 °C as a back up. RNA was eluted from the Mini Spin column with 14 uL RNase-free water (Qiagen) supplied with the RNeasy Micro kit.

### Bulk TCR repertoire profiling

For Bulk TCRα and TCRβ libraries preparation, human TCRαβ RNA Profiling Kit (MiLaboratories/Miltenyi Biotec) was used according to the manufacturer protocol (https://milaboratories.com/assets/docs/HUMAN_TCR_RNA_MULTIPLEX_KIT_Manual_1_12.pdf).

First cDNA strand was synthesized using SuperScript III reverse transcriptase (ThermoFisher Scientific, Invitrogen), 10 uL of bulk RNA were used per synthesis. To preserve RNA from degradation RNase inhibitor RNasin (Promega) was used in reverse transcription (RT). During RT each cDNA molecule was barcoded with a unique molecular identifier (UMI). These UMIs were used in subsequent bioinformatic analysis to remove PCR and sequencing errors from the data. After reverse transcription cDNA products were purified using AMPure XP Beads (Beckman Coulter) according to the manufacturer protocol. Purified TCR cDNA product was split into two equal parts for independent amplification of TCRα and TCRβ libraries and amplified in two-steps PCR amplification using Qiagen Multiplex PCR Plus kit (Qiagen). Number of PCR cycles between donors and experiments were adjusted according to the initial amount of input material (cells). For the first amplification we used 20-21 PCR cycles and for the second PCR – 12-14 cycles. During second PCR amplification, each cDNA TCRα or TCRβ library was indexed with IDT for Illumina Nextera Unique Dual Indexing primers (Illumina). Quality of obtained libraries were confirmed by capillary electrophoresis with 4150 TapeStation and D1000 or D5000 reagents (Agilent Technologies). TCRα and TCRβ libraries were pooled separately and purified with AMPure XP Beads according to the MiLaboratories manual. Concentration of TCRα and TCRβ pools was measured with 4150 TapeStation and D1000/D5000 reagents and Qubit High sensitivity dsDNA kit (Thermo Fisher Scientific). TCRα and TCRβ pools were mixed at equimolar ratio. Concentration of the final TCR pool was determined with TapeStation instrument and D1000/D5000 reagents and Qubit High sensitivity dsDNA kit. This pool was used for TCR repertoire sequencing.

PBMC replicates were lyzed either fresh or following a freeze/thaw cycle in RLT Buffer (QIAGEN) (**Supplementary Table 3**). Total RNA was isolated using the RNeasy Mini Kit (QIAGEN).

TCR repertoire sequencing was performed using Illumina NextSeq 550 or NextSeq 2000 platform in 150+150 paired-end mode, at least 50 reads per input T cell. Pool preparation was executed according to Illumina recommendations for amplicon-based libraries. To increase complexity of the sequenced pool spike with at least 15% of PhiX library was used. To minimize sample biases and batch effects, TCRα and TCRβ libraries from the same donor/experiment were sequenced within the same run.

### TCR-Seq data analysis for COVID-specific TCRs discovery

Raw fastq data were analyzed using MiXCR^37^ (v4.1.0) with the appropriate built-in preset (either milab-human-tcr-rna-multiplex-cdr3 or milab-human-tcr-rna-race-cdr3, the latter used only for pre-vaccination bulk PBMC TCR repertoire extraction, time point “pre-history”). Only cDNA molecules sequenced at least twice (based on UMI coverage) were considered in the generation of TCR clonotype tables (parameters Massemble.consensusAssemblerParameters.assembler.minRecordsPerConsensus=2 and MrefineTagsAndSort.parameters.postFilter=null in MiXCR). TCR clonotype tables were additionally filtered to include only productive CDR3β amino acid sequences starting with the conserved cysteine and ending with the conserved phenylalanine. The resulting TCRβ repertoires from bulk PBMCs were all normalized to the same depth by randomly sampling 45,000 unique cDNA molecules from each sample (ds45k dataset). To complement the missing time point before the first vaccination, an artificial “pre-history” time point was generated with a pre-vaccination TCRβ repertoire composed of exactly 45,000 cDNA molecules, based on Rep-Seq data from sorted CD4^+^ and CD8^+^ T cells from the same donor (several time points in replicates, all collected in 2017). These samples were divided into two sets, each incorporating CD4^+^ and CD8^+^ TCR repertoires, and 30,000 CD4^+^ TCR cDNA molecules and 15,000 CD8^+^ TCR cDNA molecules were randomly sampled from each set. This procedure resulted in two replicates of TCRβ repertoires from CD4^+^ and CD8^+^ T cells pooled at a ratio of 2:1 (pseudo-bulk PBMCs). All table data manipulations were performed using R (v4.1.2) and tidyverse (v1.3.2).

### Longitudinal clonotype tracking

TCR clonotypes were defined as unique nucleotide TCRβ sequences in the ds45k dataset. Differentially abundant TCRs between a target time point and three relevant reference time points, where no immune challenge was documented, were identified using edgeR (v3.36.0). Biological replicates were used to fuel edgeR statistics. The exact combinations of experimental time points together with the relevant reference time points are shown on **Fig. 2b**. Clonotypes were defined as expanded at an FDR <0.05 and a log2(count_exp_/count_ref_) >1. Artificially generated pre-vaccination TCR repertoires (pre-history) were not used in the differential abundance analysis and were only adopted for the purpose of visualization.

### Screening against datasets of TCRs with known specificity

Overlaps were sought between clonotypes in ds45k dataset and the VDJdb database ^21^ or the Pogorelyy *et al*. dataset^22^, with exact TRBV and TRBJ segment matches and a maximum of one amino acid mismatch within the CDR3 region. Clonotypes specific for SARS-CoV-2, CMV, EBV and Influenza restricted by a donor-matched HLA were analyzed. For longitudinal data, clonotypes represented by at least three unique cDNA molecules at the VAX, COVID1, or COVID2 time points in the ds45k dataset were considered reliable.

### TCRβ CDR3 homology clusters identified by three methods

Clonotypes (defined by amino acid CDR3β sequence and TRBV-TRBJ segments) identified via 1) TCR discovery, 2) clonal expansion at corresponding time points, and 3) mapping against known SARS-CoV-2-specific TCRs were pulled together before clustering, linked by TRBV-TRBJ segment and CDR3β sequence identity. This set was complemented by including clonotypes from the ds45k dataset with one amino acid mismatch. Only clonotypes with the same TRBV-TRBJ combination were allowed to be clustered together For cluster visualization, each node represents one TCRβ clonotype, and each edge links nodes with one amino acid mismatch.

### Sequencing data analysis for viral and fungi experiments

Quality of the sequenced data (fastq files) was measured using FastQC (v0.11.9) and MultiQC (v1.15) software. To reconstruct bulk TCRα and TCRβ repertoires from NGS data MiXCR software ^37^(v4.5.0, MiLaboratories) was used with a dedicated preset available for the Human TCRαβ RNA Multiplex kit (https://mixcr.com/mixcr/guides/milaboratories-human-tcr-rna-multi/). Quality control of the processed samples was performed according to MiLaboratories recommendations and included checks of alignment rate, chain usage per library and UMI threshold settings. To eliminate artificial diversity (PCR and sequencing errors) from the TCR repertoire UMI threshold cut-off was automatically applied to the samples and this threshold varied from 4 to 10 depending on the donor, sequencing depth and sample.

All subsequent data analysis was performed in R (v4.1.3). To detect TCR clonotypes that reproducibly proliferated in the presence of specific peptide pools, the *tcrgrapher* R-package (https://github.com/KseniaMIPT/tcrgrapher) was used ^36^. TCRα and TCRβ clonotypes were processed separately. First, a TCRgrapherCounts object was created with the function TCRgrapher individually for each donor by uploading all TCR datasets with the corresponding metadata information – number of replicas and antigen condition (Ag condition—PepTivator or negative control). Then a clonotype table was generated with a TCRgrapherCounts (TCRgrObject) function and default setting. Basically, in the clonotype table TCR CDR3 clonotypes with the same amino acid sequences and V gene were merged together. Then the function edgeR_pipepline was applied with the basic parameters and included levels of comparisons (Ag conditions). This function performs statistical analysis based on edgeR software from the *Bioconductor* R-package (https://bioconductor.org/packages/release/bioc/html/edgeR.html) allowing identification of significantly expanded clonotypes. Each antigen condition was compared with each other separately (Ag_1_ vs Ag_2_, Ag_1_ vs Ag_3_, Ag_1_ vs Ag_n_ etc) and only clonotypes that expanded (FDR < 0.05) under a specific condition (Ag_1_) in comparison to all other individually tested conditions (Ag_2_, Ag_3_, Ag_n_) were considered as expanded. Additionally, the following post-filtration analysis was applied to the clonotypes detected by edgeR: only clonotypes detected in at least 3 out of 8 Ag-specific replicates with a frequency above 1e4 were considered as significantly expanded.

### Subset determination

To determine initial clonotype frequency and subset identity of antigen-specifically expanded clonotypes, these clonotypes were overlapped with deep bulk TCRα and TCRβ repertoires obtained for CD4^+^ and CD8^+^ subsets before culturing. Clonotypes that overlapped only with CD4^+^ pre-cultured TCR repertoire were assigned as CD4^+^ T cell expanded clonotypes. Accordingly, clonotypes that overlapped only with CD8^+^ pre-cultured TCR repertoire were assigned as CD8^+^ T cell expanded clonotypes. If expanded clonotypes overlapped between both populations, a 5-fold ratio filter by UMI counts in CD4^+^/CD8^+^ pre-cultured TCR repertoires was applied. Clonotypes below this threshold or not detected in pre-cultured repertoires at all were classified as subset not-assigned (NA).

### Pairs prediction

Pairwise comparisons were conducted between expanded TCRα and TCRβ clonotypes to predict native TCRαβ pairs. Only TCRαβ pairs where both clonotypes were consistently present or absent in the 8 Ag-specific replicates were processed. Other candidates with discrepancy in at least one replicate (TCRα was detected but TCRβ was not present or vice versa) were excluded from further analysis. For the remaining pair candidates Pearson correlation coefficient was calculated based on the frequencies of these clonotypes across replicates. TCRαβ clonotypes with R > 0.95 and FDR < 0.05 (by Benjamini-Hochberg) were considered as reliably paired.

### Overlap with VDJdb

To confirm antigen specificity, expanded clonotypes were overlapped with VDJdb (05.06.2024). Only clonotypes with exact match by CDR3aa sequence, V-gene segment usage, and restricted by donor-specific HLA alleles were included in the analysis.

### scRNA-Seq and scTCR-Seq library preparation and sequencing

For mapping functionality of SARS-CoV-2-responsive T cell clones, frozen PBMCs from D11 (COVID1 and COVID2 samples) were thawed according to 10x Genomics recommendations and rested for 1 h in complete RPMI (RPMI 1640 + 10% autologous serum) at 37°C in the presence of 5% CO_2_. Cells were then stained with anti-CD4–Alexa Fluor 647 (clone RPA-T4, BioLegend), anti-CD8–PerCP (clone SK1, BioLegend), anti-CD19–FITC (clone J3-119, Beckman Coulter), anti-CD27–eFluor 780 (clone LG.7F9, Thermo Fisher Scientific), anti-CD45RA–eFluor 450 (clone HI100, Thermo Fisher Scientific), and 7-AAD (Thermo Fisher Scientific). Effector/memory CD4^+^ T cells gated as CD4^+^ CD8^−^ CD19^−^ after excluding CD27^+^ CD45RA^+^ events were sorted into complete RPMI using a FACSAria III (BD Biosciences). Dead cells were excluded based on morphology and staining with 7-AAD. Sorted effector/memory CD4^+^ T cells were washed twice in phosphate-buffered saline containing 0.04% bovine serum albumin and loaded onto a Next GEM Chip (10x Genomics) in one replicate with a target count of 10,000 cells for the COVID1 sample and in two replicates with a target count of 10,000 cells for the COVID2 sample. The emulsion from the COVID1 sample was divided into two parts, resulting in the generation of two technical replicates, each originating from approximately 5,000 cells. Samples were prepared using a Chromium Next GEM Single Cell V(D)J Reagent Kit v1.1 (10x Genomics). Pooled samples were sequenced with a coverage of 16,000 reads per input cell for scRNA-Seq and 5,000 reads per input cell for scTCR-Seq on a NextSeq 550 System (Illumina).

Fresh PBMCs from D01, D04, D05 were stained with anti-CCR7–PE-Cy7 (clone 3D12, BD Biosciences), anti-CD3–APC-Fire750 (clone SK7, BioLegend), anti-CD4–PE-Cy5.5 (clone S3.5, Thermo Fisher Scientific), anti-CD14–V500 (clone M5E2, BD Biosciences), anti-CD19–V500 (clone HIB19, BD Biosciences), anti-CD45RA–PE-Cy5 (clone HI100, BioLegend), and LIVE/DEAD Fixable Aqua (Thermo Fisher Scientific). Viable effector/memory CD4^+^ T cells gated as CD3^+^ CD4^+^ CD14^−^ CD19^−^ after excluding CCR7^+^ CD45RA^+^ events were sorted in two replicates for D01 and without replicates for D04 and D05 using a custom-modified FACSAria II (BD Biosciences) and loaded onto a Chromium Controller (10x Genomics). Samples were prepared using a Chromium Next GEM Single Cell 5’ Reagent Kit v2 (10x Genomics). Pooled samples were sequenced with a coverage of 100,000 reads per input cell for scRNA-Seq and 25,000 reads per input cell for scTCR-Seq on a NovaSeq 6000 System with an S4 Flow Cell (Illumina).

For IVS-selected and expanded T cells, first 2 replicates of CD137-positive expanded samples were pooled for each antigen (BZLF1 and SARS-COV-2) separately. Then, samples were prepared using a Chromium Next GEM Single Cell 5’ Reagent Kit v2 (10x Genomics). Only V(D)J-enriched libraries were sequenced, with the coverage of 5000 reads per input cell, on the NextSeq2000 system with a p1 cartridge (Illumina).

### scRNA-Seq and scTCR-Seq data analysis

Raw scRNA-Seq and scTCR-seq fastq files were processed using the count and vdj pipelines in cellranger (v6.1.2). Filtered gene expression matrices were uploaded into the Seurat R package (v4.2.0)^33^. TCR data were processed using the CellaRepertorium R package (v1.4.0). The most abundant TCR chain was used for cells with two relevant transcripts (TCRα or TCRβ). Publicly available data were downloaded via E-MTAB-10026 and converted to a Seurat object. CD4^+^ T cell clusters were chosen based on the average cluster expression of *CD4* and *CD3E*. The public dataset was then split by sample origin. CITE-Seq ADT counts were normalized using the CLR method. Additional filtering steps were performed on individual datasets, including the elimination of outlier clusters with low UMI counts and/or high percentages of mitochondrial genes and cells expressing *CD8A* and *CD8B*. The stress score was calculated for each cell using AddModuleScore() function in Seurat and included genes upregulated as a consequence of the dissociation procedure^38^. Integration was carried out using the Seurat reference-based reciprocal PCA protocol, with the largest 3’ and 5’ datasets chosen as references to account for differences in methodology^33^. The percentage of mitochondrial genes and the stress gene signature were regressed out of individual and integrated datasets. All TCR and IG genes were removed from variable features used in the PCA and from the anchor features used for integration. The number of dimensions used for running the UMAP algorithm was 25.

### Reference mapping

D11_COVID1, D11_COVID2_rep1, and D11_COVID2_rep2 samples were mapped to the integrated reference dataset^24^ using Seurat Reference Mapping procedure with default parameters^33^. Plots were designed to highlight amino acid-defined TCRβ clonotypes with specificity for SARS-CoV-2.

## Notes

### Competing Interest Statement

Egorov E.S., Martinez Carrera L.A., Jelveh N., Bisdorf K., Matzke S.K., Khorkova S., Bosio A., Hardt O., Serebrovskaya E.O. are employed by Miltenyi Biotec B.V. & Co. KG, Bergisch Gladbach, Germany

