## Supplementary data for "Discovery of rare antigen-specific TCRs via replicate profiling"

**Supplementary Note 1**. Patient timeline and sample collection.

Male donor D11 was born in 1978, had no chronic diseases at the start of the study. Vaccinated with the two-dose Sputnik V in Dec2020/Jan2021. HLA genotype: A*03:01:01, A*25:01:01, B*15:01:01, B*40:02:01, B*40:303, C*03:04:01, C*03:03:01, C*03:20N, C*03:227, DQB1*06:02:01, DQB1*05:01:01, DRB1*01:01:01, DRB1*15:01:01. D11 got two registered, PCR-confirmed SARS-CoV2 infections, in April 2021 (moderate illness, strain B.1.1.7 alpha, confirmed by virus sequencing), and January 2022 (mild illness, supposedly strain omicron, based on the current epidemiology). At multiple time points, PBMC were isolated in replicates. Performed analyses included: 1) IFN-γ ELISpot for SARS-CoV2 antigens; 2) Deep TCRβ repertoire profiling (TCR-Seq) with HUMAN TCR RNA MULTIPLEX kit from MiLaboratories Inc; 3) scRNA-Seq and scTCR-Seq on the time points right after the first and second infections. 4) TCR discovery assay on the time point right after infection. See **Supplementary Fig**. **1a** for the patient timeline. IFN-γ ELISpot showed T cell response to Miltenyi spike protein (S) peptivator after Sputnik V vaccination, and to both spike and nucleoprotein (N) at both time points of SARS-CoV2 infections (**Supplementary Fig**. **1b**).

**Supplementary Table 3**. Number of predicted pairs (n=201) in 5 individuals stimulated with different PepTivators.

| Donor / Peptivator | CMV | EBV | SARS-CoV-2 | MP65 | CEF |
|---|---|---|---|---|---|
| **B26** | 9 | 1 | - | - | - |
| **C76** | 42 | - | 56 | - | - |
| **C34** | - | 3 | 14 | - | 10 |
| **C26** | - | 11 | - | 10 | - |
| **C97** | - | 37 | - | 8 | - |

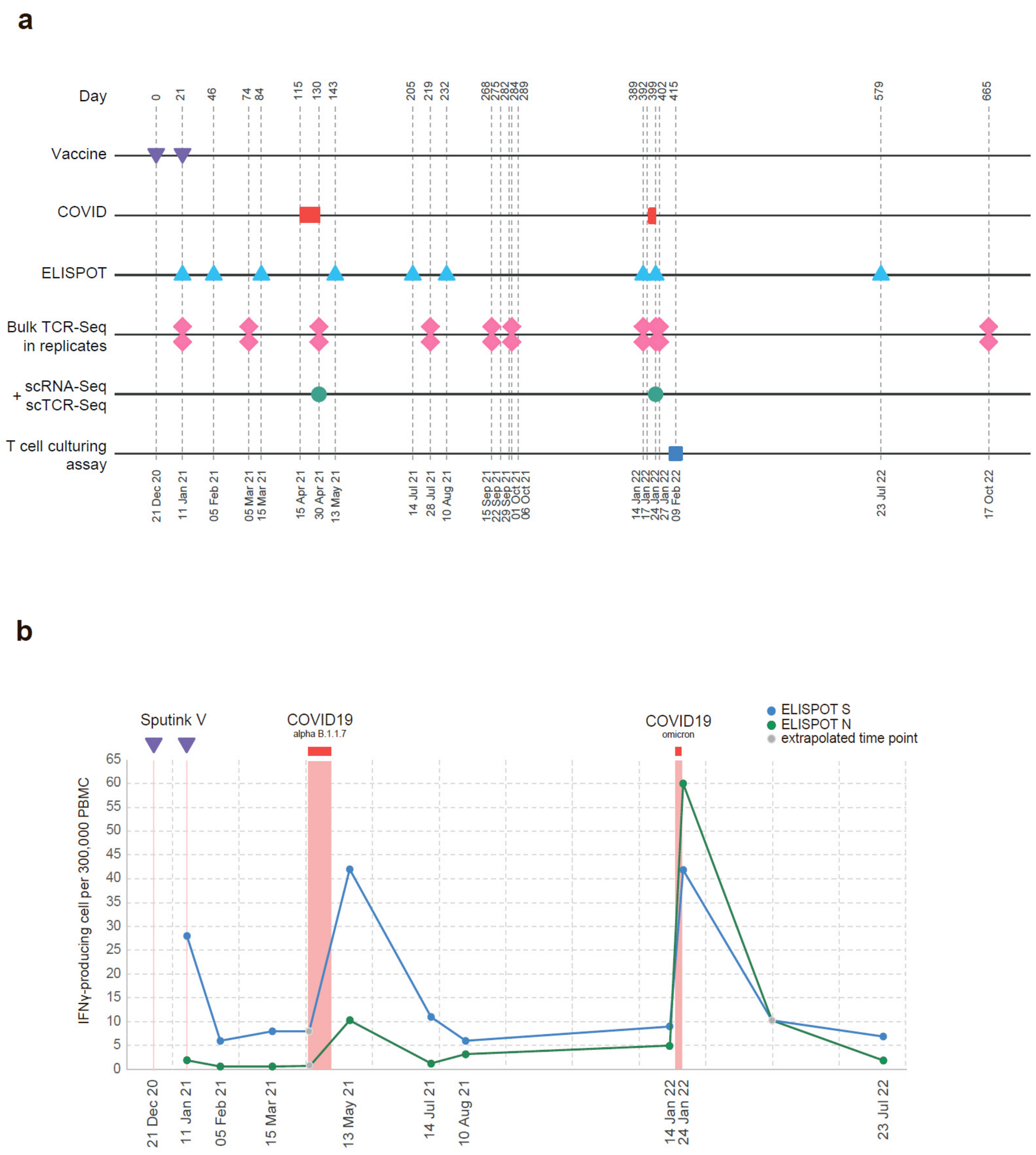


**Supplementary Figure 1. D11 donor timeline**. **a.** Experimental design. D11 was tracked for two years. Two shots of adenovirus-based vaccine Sputnik V and two break-through COVID infections are shown. ELISPOT: IFNγ ELISPOT assay on PBMC; Bulk TCR-Seq was performed from PBMC; scRNA-Seq and scTCR-Seq were performed from sorted Effector/Memory CD4 T cells. **b**. IFNγ ELISPOT assay results. S- and N-peptivator peptide pools were used as antigens. Sputnik V contains full-length Spike protein-encoding sequence; Gray dots - extrapolated data points.

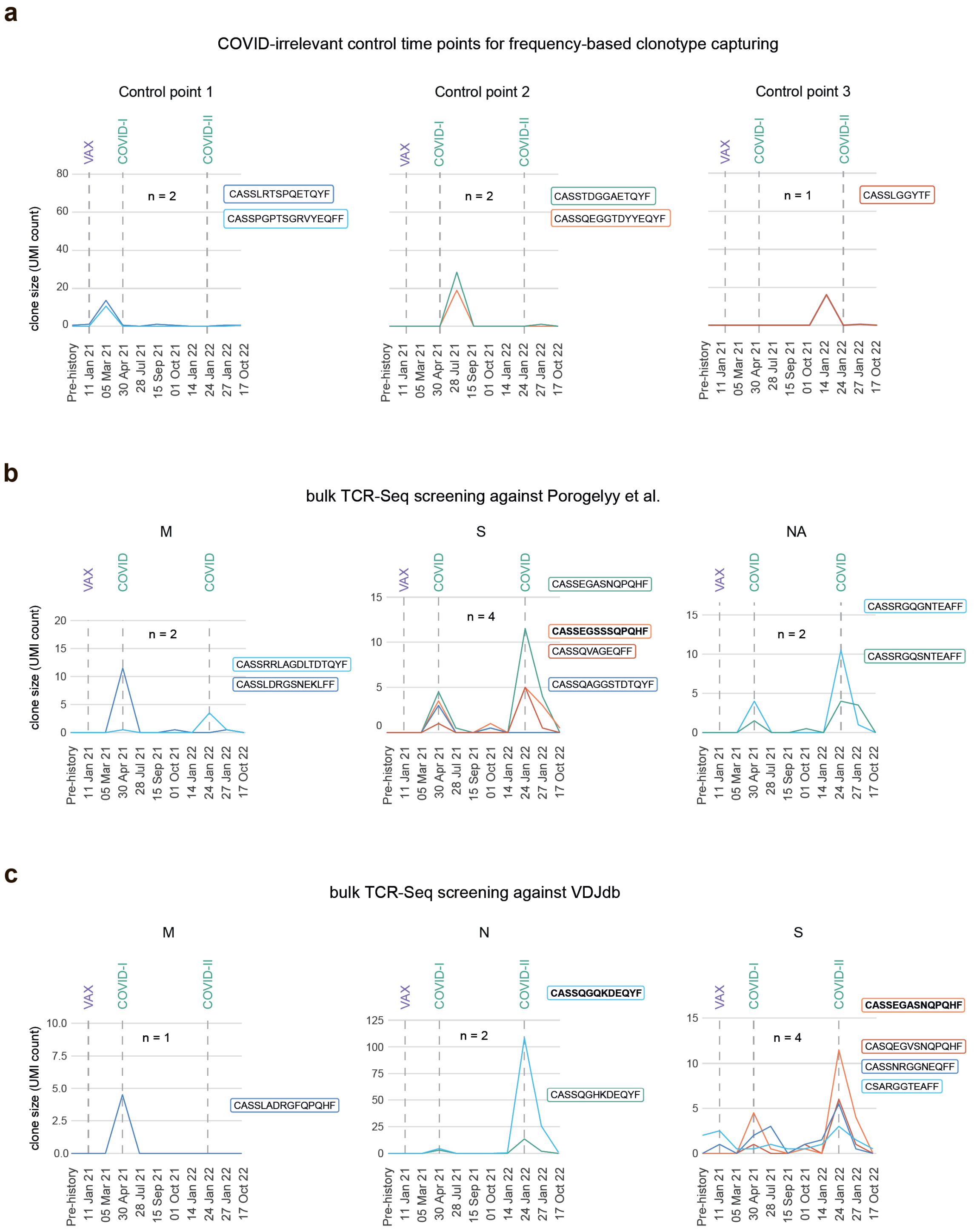


**Supplementary Figure 2. COVID-specific TCR clonotypes identified based on overlap with databases. a.** Longitudinal tracking of TCRβ clonotypes (CDR3nt + V +J) identified based on the expansion of frequency at control time points (controls for false discovery) **b**,**c.** Longitudinal tracking of TCRβ clonotypes (CDR3aa + V + J) overlapping with published data[1] (b), or VDJdb[2, 3] (c). Up to one amino acid mismatch is allowed. One clone permanently present at high frequency in the peripheral blood of the donor during the whole observational period starting from 2017, and assigned as Spike-specific in VDJdb, was removed from the analysis. Each PBMC TCR-Seq library is downsampled to 45,000 TCRβ encoding cDNA molecules (UMI). TCRβ CDR3 amino acid sequences are shown.

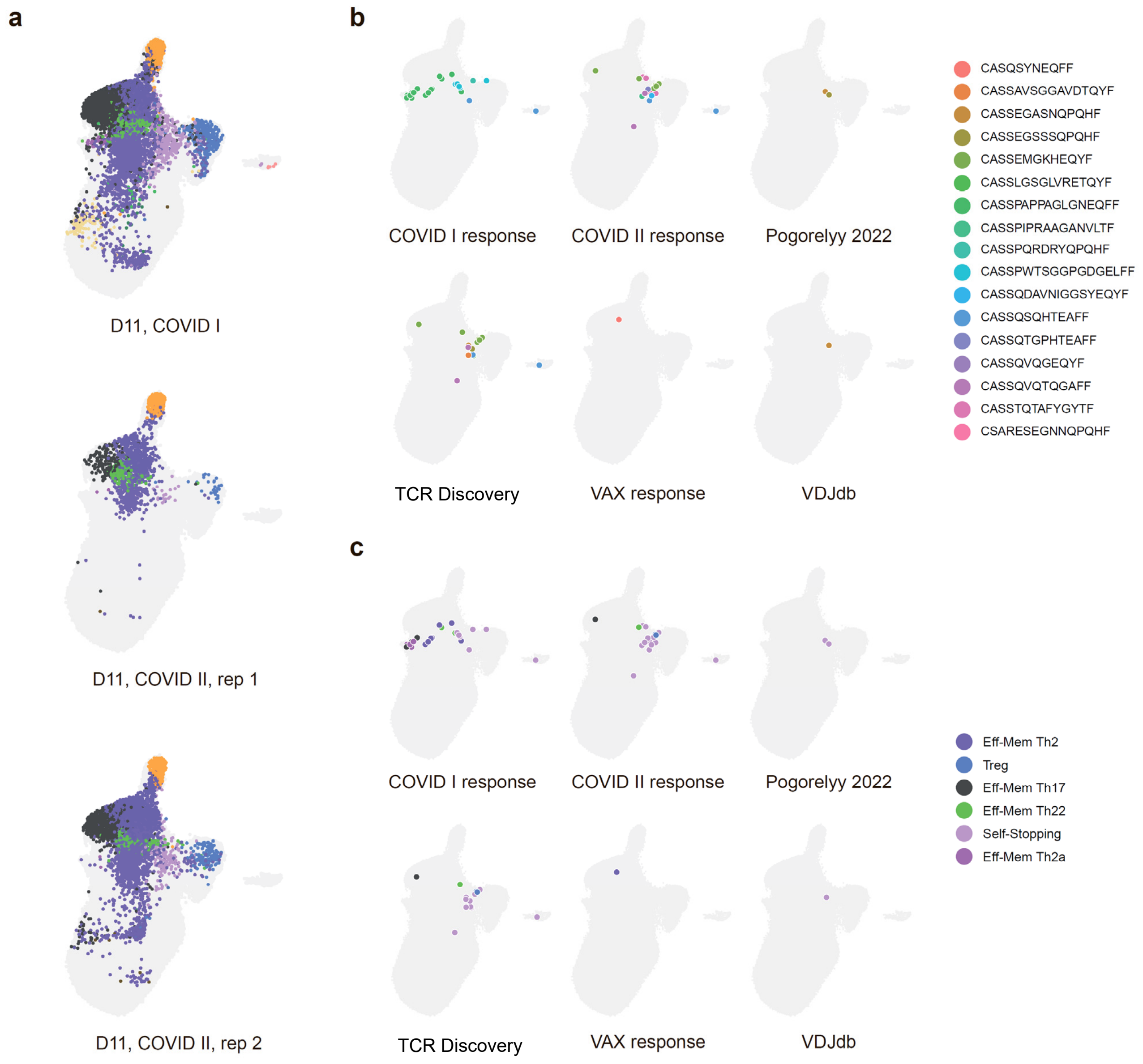


**Supplementary Figure 3. Mapping to the reference dataset shows functional clusters of SARS-CoV-2-specific CD4+ T cells. a**. UMAP plots showing cells from the reference dataset (in gray) and mapped effector-memory enriched CD4+ T cells from D11 donor. **b,c.** Positioning of SARS-CoV-2-specific TCRβ CDR3 clonotypes. SARS-CoV-2-specific cells are colored based on clonotype (b), and scRNA-Seq cluster (c).

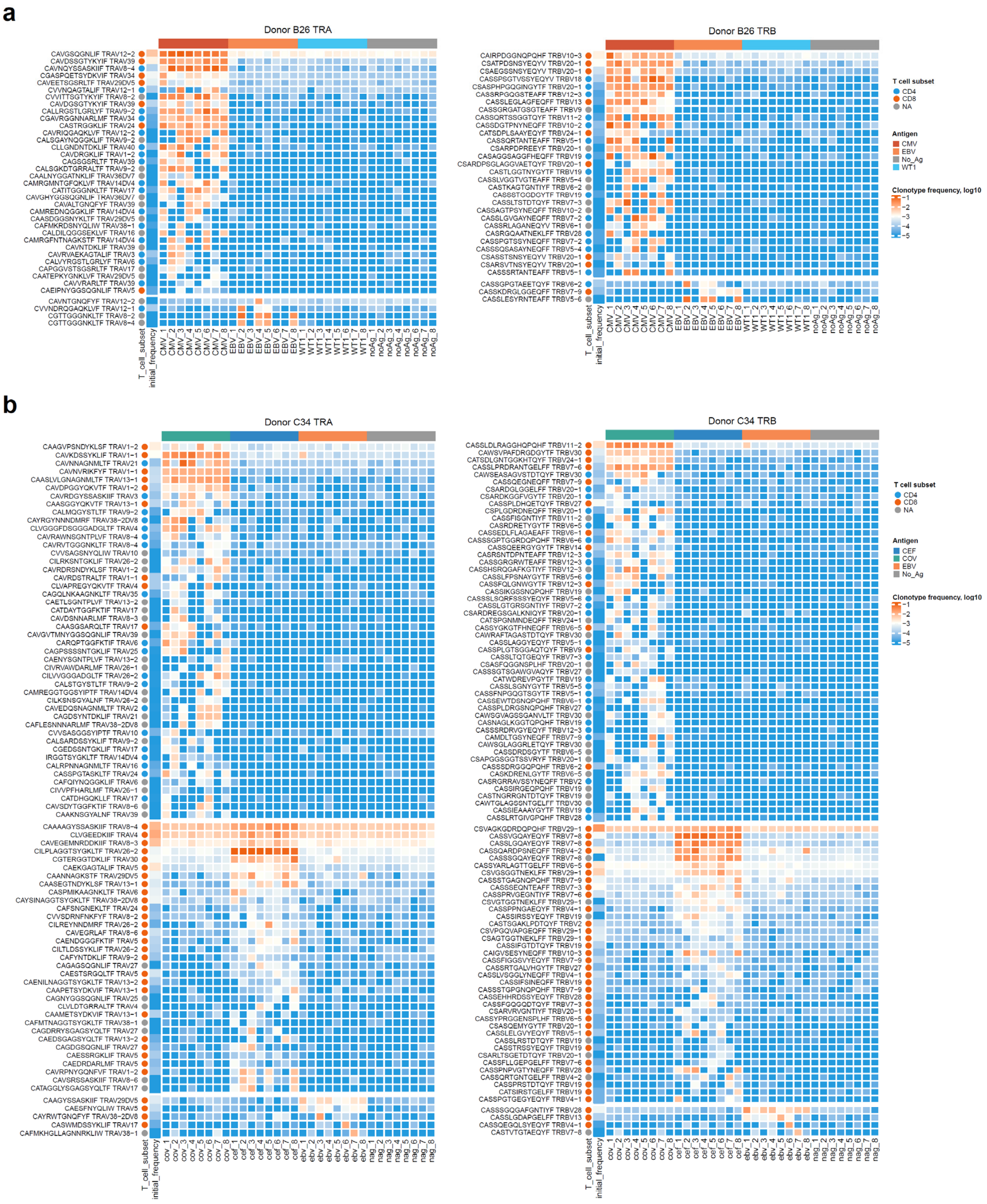


**Supplementary Figure 4.** Viral-specific TCRs. **a,b.** Frequency of TCRα and TCRβ clonotypes specifically expanded in response to PepTivators CMV pp65 and EBV BZLF1 for Donor B26 (a), and PepTivators SARS-CoV-2 Prot_S, CEF MHC Class I Plus, EBV BZLF1 for Donor C34 (b).

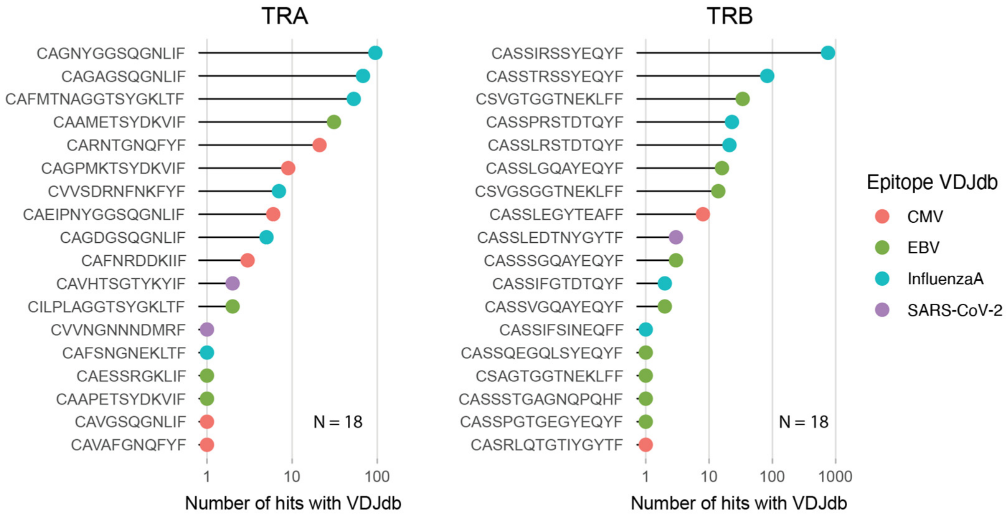


**Supplementary Figure 5**. Viral-specific CD8 TCRα (N = 18) and TCRβ (N = 18) clonotypes detected by TCR-Disco that matched entries in the VDJdb database. Only clonotypes with fully matched CDR3 amino acid sequences, V gene segments, and MHC class I context were considered valid matches.

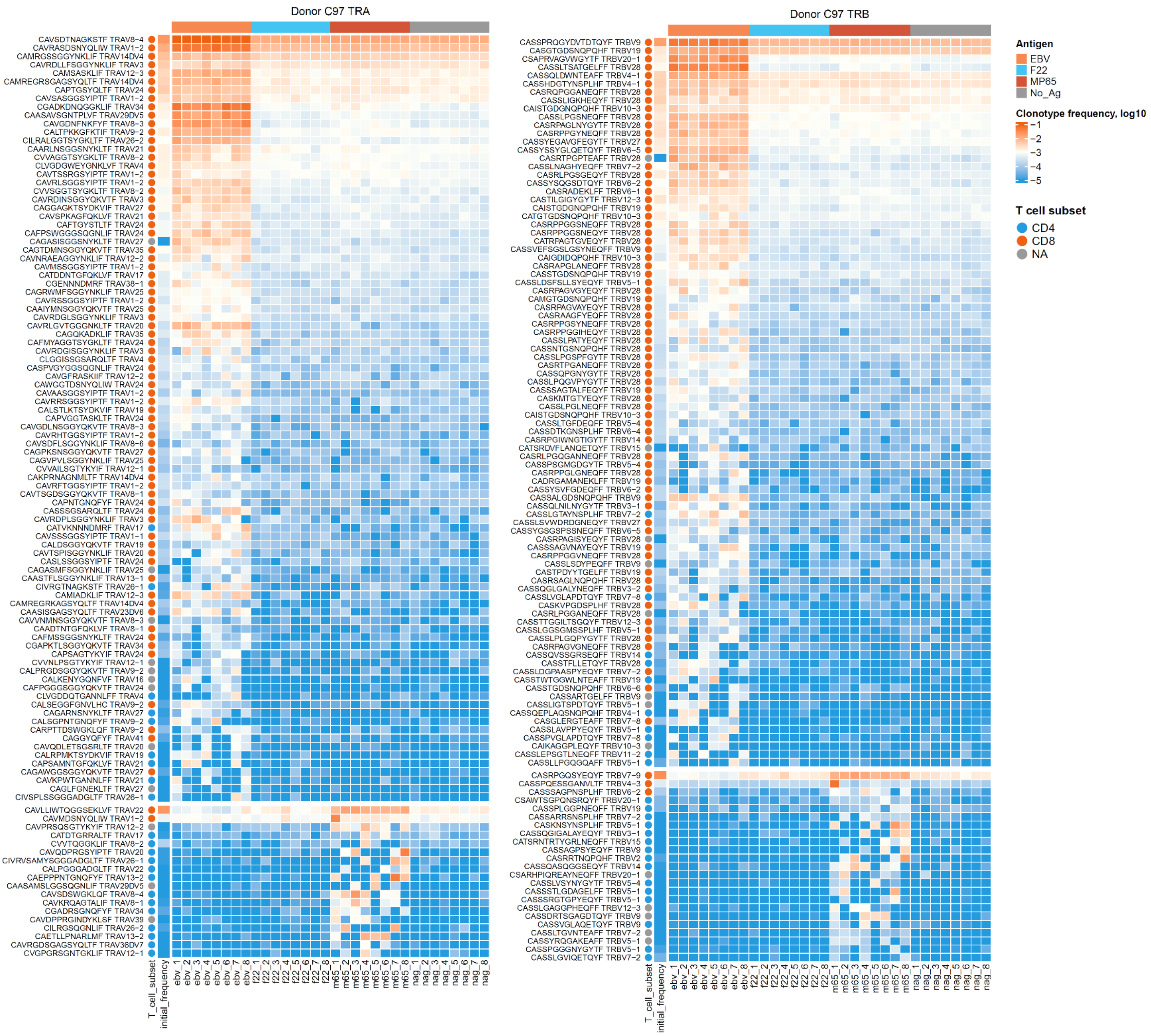


**Supplementary Figure 6. Fungi-specific TCRs.** Frequency of TCRα and TCRβ clonotypes specifically expanded in response to EBV Consensus and C. albicans MP65 PepTivators for donor C97.
